# A spatiotemporally resolved GPCR interactome reveals novel mediators of receptor agonism

**DOI:** 10.1101/2024.06.14.599010

**Authors:** Maria M. Shchepinova, Rachel Richardson, Jack W. Houghton, Abigail R. Walker, Mohammed A. Safar, Daniel Conole, Aylin C. Hanyaloglu, Edward W. Tate

## Abstract

Cellular signaling by membrane G protein-coupled receptors (GPCRs) is orchestrated by a complex and diverse array of mechanisms. The dynamics of a GPCR interactome as it evolves over time and space in response to an agonist can offer a unique window on pleiotropic signaling decoding and functional selectivity at a cellular level. In this study, we employed proximity-based APEX2 proteomics to interrogate the interaction network of the GPCR for luteinizing hormone (LHR) on a minute-to-minute timescale. We developed an analytical approach integrating quantitative multiplexed proteomics and temporal reference profiles, providing a platform to identify the proteomic environment of APEX2-tagged LHR at the nanometer scale. LHR activity is exquisitely regulated at a spatial level, leading to identification of novel putative interactors including the Ras-related GTPase RAP2B that modulate both receptor signaling and post-endocytic trafficking, and providing a resource for spatiotemporal nanodomain mapping of LHR interactors across subcellular compartments.

**Significance:** G-protein coupled receptors (GPCRs) are the largest family of membrane proteins in the human body. GPCRs have established themselves as key drug targets, playing central roles in health and disease. The activity of this superfamily of receptors is highly dynamic, integrating diverse signaling pathways over time and across multiple subcellular compartments. Quantitative proximity proteomics using engineered ascorbate peroxidase (APEX2) has emerged as a powerful tool to map GPCR-protein interaction landscapes in both time and space. However, resolving these networks at a sub-minute timescale remains challenging, particularly for poorly characterized GPCRs with limited pharmacological tools or complex trafficking patterns. In this study, we applied APEX2 proximity proteomics to resolve at nanoscale the agonist (LH)-induced interactome of the luteinizing hormone receptor (LHR), a key GPCR in human reproduction. LHR undergoes rapid endocytosis upon LH stimulation, trafficking to a poorly understood compartment termed the very early endosome (VEE), where it engages in G protein signaling and post-endocytic sorting. Harnessing the temporal profile of GIPC1 protein, a key known interactor of LHR in the VEE pathway, we developed an analytical pipeline to identify novel endocytic interactors of LHR. Using our platform, we identified RAP2B and RAB38 as potential modulators of LHR signaling and trafficking. Overall, we present an APEX2 pipeline optimized for interrogating complex datasets, particularly for GPCRs, and potentially other membrane receptors, with limited known interactome data and minimal pharmacological tools. Our findings highlight both the complexity of GPCR interaction networks and an effective strategy to deconvolute them, providing a comprehensive resource to advance understanding of this large family of receptors.

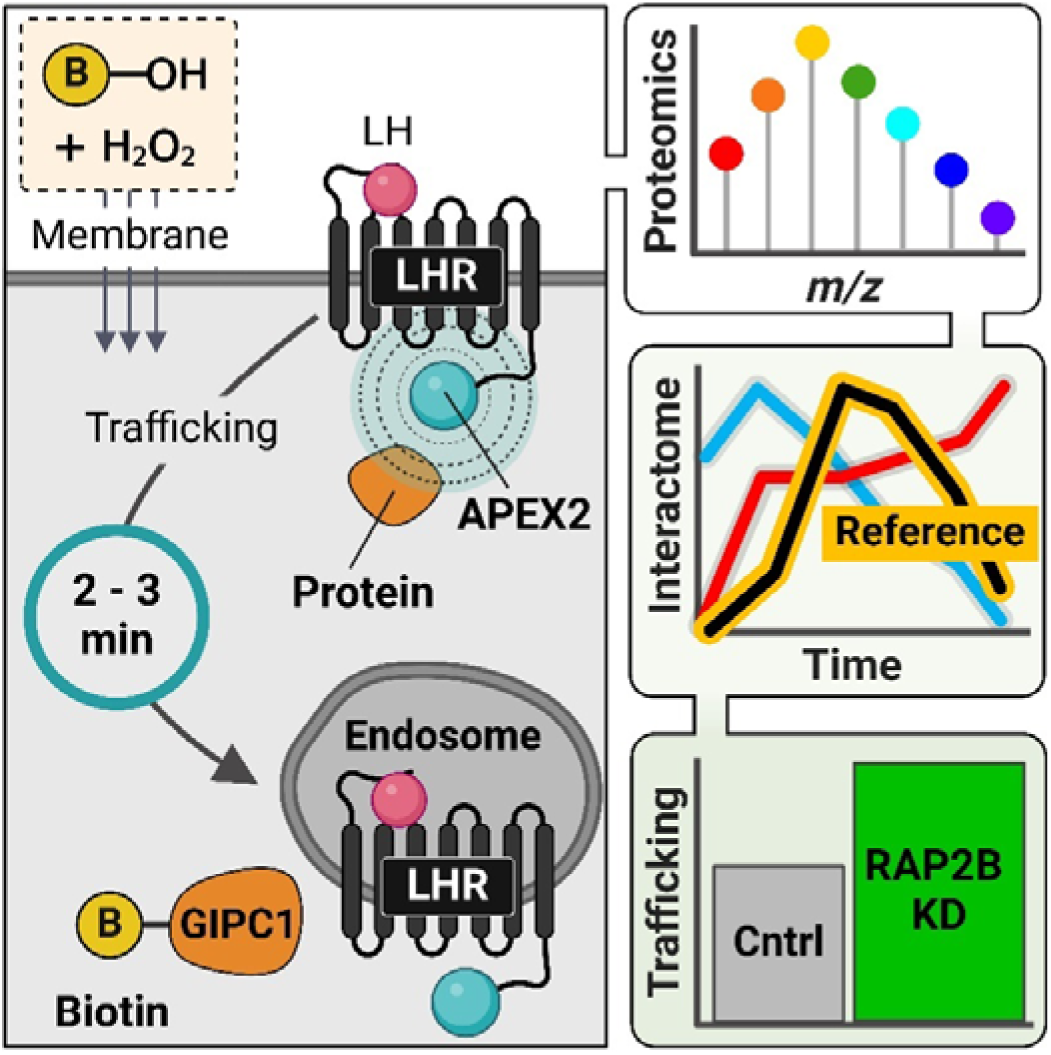

## Introduction

The G protein-coupled receptor (GPCR) superfamily plays a central role in cellular communication and signaling ^1–3^. Current models of GPCR signaling have evolved beyond a single GPCR activating a specific heterotrimeric G protein pathway at the cell surface to encompass dynamic multidimensional signaling hubs ^4^. These include the ability of GPCRs to activate multiple G proteins, associate with an array of adaptor proteins ^5^ and form oligomeric complexes ^6,7^. Furthermore, GPCRs assemble distinct signaling hubs in varied subcellular compartments in addition to the plasma membrane, including endosomes ^8^, the nucleus ^9^, Golgi and mitochondria ^4,10^. Collectively, this diversity of mechanisms offers multiple layers to regulate functional selectivity for a specific GPCR ligand ^4^. To understand the relevance of GPCR signaling in health and disease, we must develop methods to learn how such complexity is decoded by cells to direct downstream functions.

The luteinizing hormone receptor (LHR) is an example of a GPCR that exhibits cell signaling pleiotropy (the ability of the receptor to produce several different signaling outputs in response to ligand binding). This pleiotropy is thought to mediate its diverse roles in reproduction and pregnancy, harnessing poorly understood mechanisms to regulate signaling activated by its endogenous ligand luteinizing hormone (LH) at a spatiotemporal level (**Figure 1A**) ^11–13^. Whilst LHR is primarily coupled to the heterotrimeric G protein G_α_s, thereby increasing levels of the second messenger cyclic AMP (cAMP), under conditions of high LHR expression and/or LH levels observed in ovaries, it may also activate G_α_q/11 signaling pathways, leading to activation of phospholipase C-β (PLC-β) ^14–16^. Activated LHR also robustly recruits canonical GPCR adaptor proteins β-arrestin-1 and -2 for rapid G protein desensitization, internalization from the plasma membrane to endosomes and activation of additional signaling pathways, such as mitogen activated protein kinases (MAPKs) ^12,17,18^. Indeed, the location or spatial control of LHR is critical for shaping LHR activity, whereby receptor internalization is essential for its acute G_α_s-cAMP signaling ^19^. The LHR initiates this G protein signaling via receptor internalization and trafficking to a unique endosomal compartment termed the very early endosome (VEE), which plays a key role in sorting receptors to post-endocytic trafficking pathways (**Figure 1A**) ^19,20^. In contrast to the early endosome, the VEE is of smaller size, and lacks classical early endosomal markers such as EEA1, phosphatidylinositol 3-phosphate (PI3P) and Rab5 ^20^. LHR is a paradigm example of a GPCR regulated by the VEE, which is a platform for acute intracellular G protein signaling, sustained MAPK signaling mediated by interactions with the PDZ domain protein GIPC1 (G_α_i-interacting protein C-terminus 1) during ligand-induced endocytosis, and rapid recycling to the plasma membrane, driven by the protein APPL1 (Adaptor protein containing PH domain, PTB domain and Leucine zipper motif), which also functions in the VEE to negatively regulate LHR G protein signaling via distinct mechanisms ^19,21^. APPL1, the only adaptor protein previously shown to localize to VEE, marks only a subpopulation of LHR-containing VEEs ^19^, and the protein machinery driving LHR activity through these critical sorting and signaling compartments remain poorly understood.

**Figure 1.**
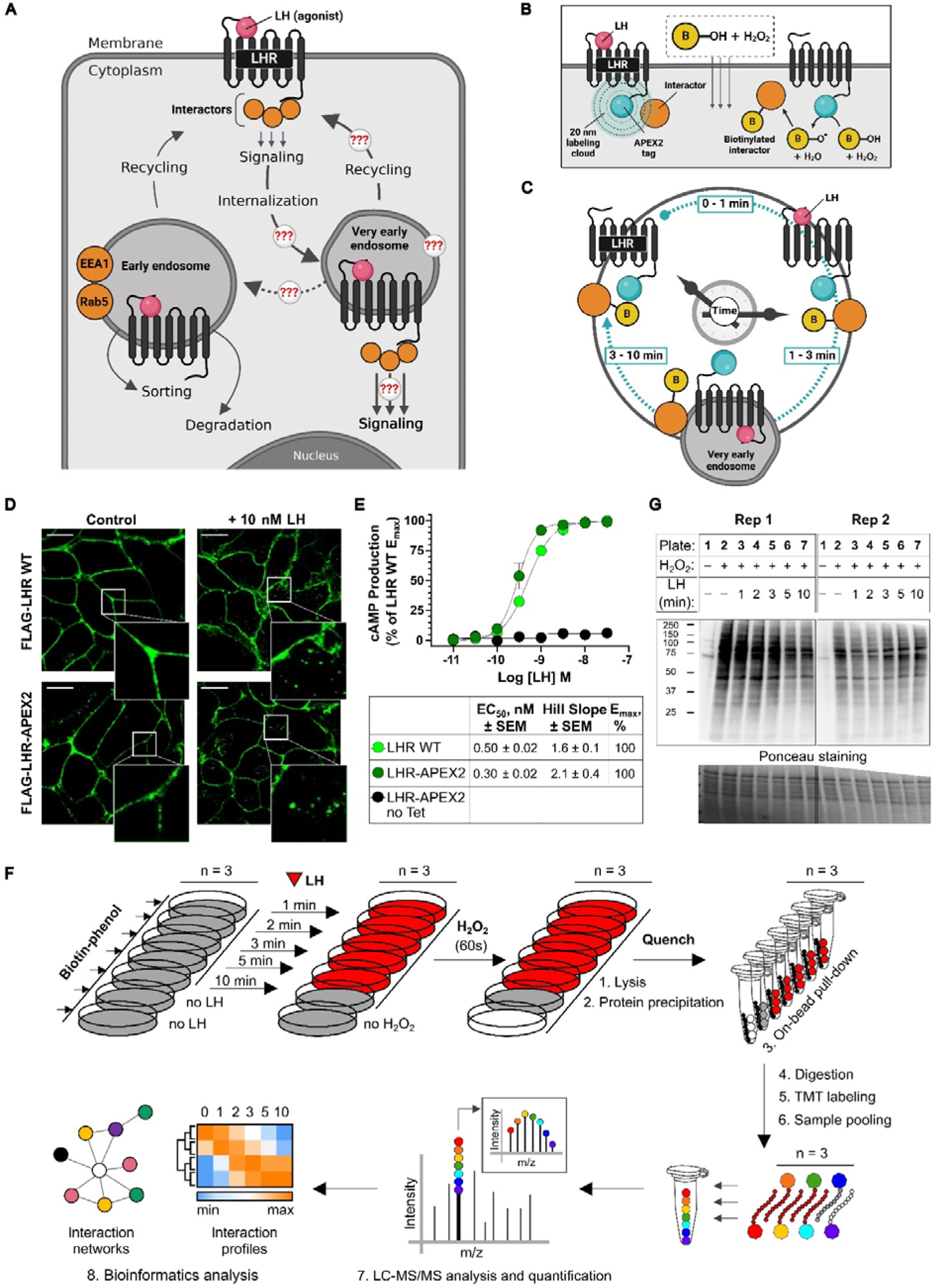
Establishing an LHR-APEX2 proximity labeling platform. **A**. Model summarizing current understanding of signaling and trafficking pathways of LHR activated by LH agonist. Upon activation, LHR couples to interactors that mediate its downstream effects including cAMP signaling, desensitization and internalization. Following internalization, LHR is trafficked to very early endosomes (VEE), smaller in size than early endosomes and lacking classical endosomal markers EEA1 and Rab5. From the VEE, LHR elicits downstream signaling cascades, including cAMP generation. LHR is then primarily rapidly recycled back intact to the plasma membrane, with a minor proportion sorted towards the EE and subsequently degradation ^11^. However, the dynamic LHR interactome during endocytic trafficking to and from VEEs is currently unknown. **B**. To enable APEX2 proximity labeling, the APEX2 enzyme ^22^ is fused to the C-terminus of LHR where it can catalyze the formation of biotin-phenoxy radicals from biotin-phenol in the presence of hydrogen peroxide. These radicals persist only in close proximity (<20 nm) to LHR resulting in rapid biotinylation of nearby protein interactors, which can then be enriched, identified and quantified by quantitative mass spectrometry-based proteomics ^23^. See also **Figure S1A**. **C**. Timeline diagram of APEX2 proximity labeling along LHR endocytic signaling and trafficking route represented by its agonist-dependent functional ‘clock’ ranging from 1 up to 10 minutes post agonist addition. **D**. Validation of LHR-APEX2 expression and trafficking by confocal microscopy. FLAG-LHR WT HEK293 cells and Flp-In T-REx 293 FLAG-LHR-APEX2 cells were labeled with anti-FLAG antibody and non-treated (control) or treated with 10 nM LH for 10 min. Cells were fixed and stained with a secondary AlexaFluor 488 (AF488)-conjugated antibody. Shown is a representative frame from imaging of cells via confocal microscopy with a zoom-in inset (bottom right). Scale bar = 5 μm. Both untagged and APEX2-tagged LHR constructs exhibit comparable plasma membrane expression and LH-induced internalization into VEEs. See also **Figure S1B**. **E**. Validation of LHR-APEX2 signaling by TR-FRET cAMP assay normalized to the LHR WT E_max_. Dose-response cAMP curves for the FLAG-LHR WT HEK293 cells and tetracycline-induced Flp-In T-REx 293 LHR-APEX2 cells activated by various concentrations of LH produce comparable EC_50_ and E_max_ values. Non-induced cells do not respond to LH treatment. Data represent mean ± SEM (n = 3). **F**. APEX2 proximity labeling and mass spectrometry analysis workflow. Flp-In T-REx 293 LHR-APEX2 cells were pre-incubated with biotin-phenol, treated with LH for different time points, activated with hydrogen peroxide and quenched. Biotinylated proteins were pulled-down on streptavidin-coated beads, digested, labeled with the tandem isobaric mass tag (TMT) kit, analyzed and quantified by LC-MS/MS, followed by bioinformatics analysis. **G**. Biotinylated proteins analyzed by neutravidin-HRP western blot; increased biotinylation was observed for hydrogen peroxide-treated as opposed to non-treated samples. See also **Figure S1C**.

Proximity labeling-enabled proteomics of interacting proteins (interactomics) is an emerging and powerful approach to interrogate protein interaction networks at both the spatial and temporal level, directly in intact cells ^24–29^. The approach applies proximity-based biotinylation through a fusion of the protein of interest with APEX2, an engineered ascorbate peroxidase domain ^22^, coupled to enrichment and quantitative mass spectrometry proteomics. APEX2 proximity proteomics was first applied to GPCRs to identify known and novel interactors of β_2_-adrenergic receptor ^30,31^, angiotensin II type 1 receptor ^30^, δ-opioid receptor ^31^ and μ-opioid receptor ^32^. Rapid labeling within 30 seconds over a distance of up to 20 nm from the GPCR-APEX2 fusion offers a particular advantage for the analysis of GPCR signaling interactome, where receptor activation, internalization and recycling typically occurs on a timeframe of seconds to minutes ^33^. Here, we employed LHR-APEX2 fusions to interrogate the ligand-induced LHR interaction network, with minute to minute temporal resolution. Harnessing known time-dependent LHR-protein associations through a multi-layered quantitative interactome analysis pipeline, we established the first approach to spatiotemporally identify and characterize the LHR interactome during receptor activation in intact cells and identify multiple novel putative LHR interactors which may direct very early endosomal sorting and signaling. Taken together, this study provides a roadmap for analysis and functional interrogation of receptor-mediated quantitative proximity labeling proteomics datasets.

## Results

### Establishing an LHR-APEX2 proximity labeling platform

To validate our system, we set out to apply an LHR proximity labeling platform (**Figure 1B**) to identify the LH (agonist)-dependent LHR interactome over a 1-10 minute time window (**Figure 1C**), with the aim to capture well-established key steps in receptor signal activation, internalization, and subsequent trafficking, based on the current knowledge of LHR activity profiles (**Figure 1A**) ^19,20^. We stably expressed LHR C-terminally tagged with APEX2 ^22^ in tetracycline-inducible Flp-In T-REx 293 cells ^34^ (**Figure S1A**), and went on to confirm that this novel fusion preserved the expected expression, internalization and recycling behavior of LHR ^19,20^. LHR-APEX2 exhibited plasma membrane expression and LH-induced internalization to small endosomes characteristic of VEEs, in a manner identical to cells stably expressing N-terminally tagged FLAG-LHR without APEX2 fusion (**Figure 1D**), and it also recycled to the plasma membrane (**Figure S1B**). The ability of LHR-APEX2 to activate G_α_s-cAMP was also compared to FLAG-LHR expressing cells, and both cell lines were found to produce similar LH-mediated cAMP dose-response curves with comparable values for half maximal effect concentration (EC_50_) and maximum response (E_max_) (**Figure 1E**). Cells without tetracycline treatment did not show detectable LHR activity upon activation with LH (**Figure 1E**). Taken together, these results indicate that C-terminal APEX2 tagging did not impair protein interactions required for recycling or activation of key G protein signal pathways, and the key elements of the interactome were preserved.

To optimize the proximity labeling reaction (**Figure 1B**), Flp-In T-REx LHR-APEX2 cells were induced with tetracycline overnight, and then pre-incubated in biotin-phenol for 30 min followed by stimulation with the agonist LH for either 1, 2, 3, 5 or 10 min (**Figure 1F**). Each sample was treated with hydrogen peroxide for 60 seconds to initiate APEX2-catalyzed proximity biotinylation, followed by immediate quenching with ice-cold quenching buffer. For controls, “no hydrogen peroxide” or “no agonist” treatments were used, as in previous APEX studies ^23,30,31,35,36^.

Cells were harvested, lysed, and proteins precipitated. The levels of streptavidin-HRP, as measured by western blotting, demonstrated strong biotinylation present in all samples apart from controls lacking hydrogen peroxide treatment (**Figure 1G**), which revealed only endogenous biotinylated carboxylases at ca. 75 and 130 kDa, as expected ^23,37^. These biotinylated proteins could be successfully pulled-down on streptavidin-coated agarose beads (**Figure S1C**).

For subsequent proteomics analysis (**Figure 1F**), “no hydrogen peroxide” and “no agonist treatment” controls were used, and proximity labeling was carried out in three independent replicates.

### Quantitative proximity labeling enables dynamic activation-dependent LHR interactomics

To identify and quantify LH-responsive biotinylated proteins, we established a multi-step proteomics pipeline (**Figure 1F**). Enriched biotinylated proteins were digested on-bead, peptides quantified, concentrated and labeled with 7-plex isobaric tandem mass tags (TMT); peptides were combined per replicate, fractionated by alkaline reverse phase LC, separated by nanoscale reverse phase LC, and analyzed by tandem mass spectrometry (MS/MS). Samples with no hydrogen peroxide treatment were included as a background control and samples with no LH treatment were used as non-stimulated LHR-APEX2 controls. Across three independent experiments, 3584 proteins were quantified in MaxQuant ^38^ (**Figure 2A**). As expected, the majority of proteins identified in no hydrogen peroxide-treated samples showed significantly lower protein abundances than hydrogen peroxide-treated samples, in agreement with western blotting data (**Figure S2A**). After log2 transformation, proteins with comparable abundances across all samples including the “no hydrogen peroxide” background control were considered non-specific and omitted from further analysis in Perseus ^39^ via the negative outlier removal step (FDR <0.05) (see **Data S1** for full quantification data tables). Next, the enriched protein abundances were normalized to LHR-APEX2 within each sample as an internal reference for labeling efficiency, protein level, pull-down efficiency and sample processing (**Figure S2B-E**). To focus analysis on the activation-dependent LHR-APEX2 interactome, proteins that either decreased or did not change in abundance following LH treatment were removed, followed by ComBat correction to remove batch effects ^40^. For that, all proteins were normalized per replicate to “no LH treatment” control and their maximal abundance values for at least one time point following receptor activation were tested for significance over “0” (“no LH treatment”) (FDR <0.05), and subsequent analyses were carried out on proteins that displayed increased enrichment, and therefore potentially increased interaction with LHR (**Figure S2F**). In total, 2439 proteins were taken forward to KEGG pathway analysis using the DAVID online platform ^41^, confirming endocytosis as the most enriched pathway, as anticipated, with 122 proteins identified (**Figure 2B**). Similarly concordant enriched pathways included tight junctions, regulation of actin cytoskeleton, small G protein signal pathways (e.g. Rap1 and Ras) and the Hippo signaling pathway.

**Figure 2.**
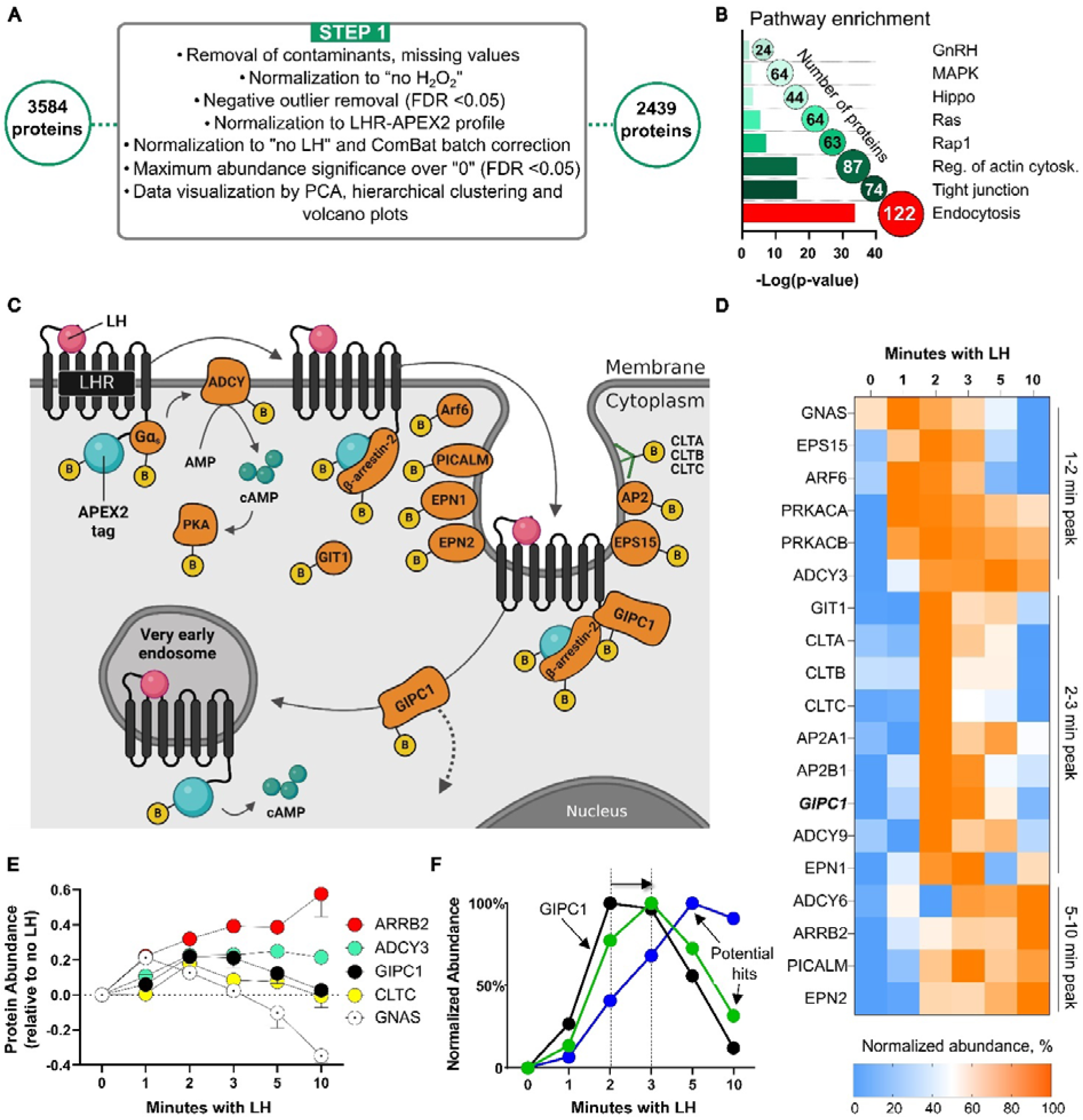
APEX2 captures known agonist-dependent LHR interactome. **A**. Protein filtration and normalization steps. For 3584 proteins found in MaxQuant ^38^, contaminants and missing values were removed in Perseus ^39^, and proteins significantly enriched over background (“no H_2_O_2_” control) were identified in a Significance A outliers test (FDR <0.05). The internal reference (LHR-APEX2 profile) was subtracted from all protein profiles, proteins were then normalized to the non-simulated (“no LH”) control, and ComBat batch corrected ^40^. Maximum abundance values across all time points per replicate were subjected to a t-test (Benjamini-Hochberg FDR <0.05) to identify significantly enriched proteins following LH treatment, resulting in 2439 proteins. See also **Figure S2** and for the list of proteins and data tables refer to **Data S1**. **B**. Selected KEGG pathways enriched in proteomic datasets according to the DAVID online analysis platform^41^. Endocytosis is the most statistically enriched pathway in the dataset, with the highest number of proteins (122) represented. **C**. Model summarizing known LHR interactors revealed by proteomic analysis. GIPC1 protein ^20,42^ must dissociate from the LHR C-terminal domain before LHR enters VEEs. **D**. Heatmap of interaction profiles for known LHR interactors. Protein abundance (log2 fold change) for each protein across different time points was normalized to the non-stimulated (“no LH”) control first, followed by normalization to its maximum and minimum enrichment signal. The profiles are ordered from early to late, according to the timepoint of their strongest interaction with LHR. For hierarchical clustering of the heatmap refer to **Figure S2G**. For other G proteins and Hippo pathway proteins found in the dataset refer to **Figures S3** and **S4**, respectively. **E**. Temporal profiles of selected known LHR interactors. Protein abundance (log2 fold change) for each protein across different time points was normalized to the non-stimulated (“no LH”) control. Data represent mean ± SEM (n = 3). **F**. Schematic illustration of potential VEE-related hit interactor temporal profiles (green and blue), set against the reference profile for GIPC1 (black). Peak profiles for potential VEE-related hits are shifted to later timepoints compared to GIPC1, which peaks at 2 min and dissociates from the LHR C-terminus before LHR enters VEEs, providing a temporal reference point for VEE entry.

### A time-resolved LHR interactome referenced to known LH-dependent LHR interactors

In order to establish reference points to identify and functionally classify novel LHR protein partners, we first analyzed the time-resolved interaction profiles of known ligand-dependent GPCR and/or LHR interactors. We found that profiles for multiple known agonist-dependent LHR interactors were consistent across three independent replicates, indicating remarkably robust reproducibility of proximity labeling. These included canonical GPCR interactors G_α_s protein (GNAS), β-arrestin-2 (ARRB2), LHR C-terminal tail interactor GIPC1 ^42^, which followed profiles based on previously reported data using orthogonal methodologies ^20,43,44^, and known GPCR signaling and trafficking network components with interaction profiles similar to those found by other GPCR-APEX2 studies ^30,31^, including adenylate cyclases (ADCY), protein kinase A (PRKACA and PRKACB), clathrin-mediated endocytosis components (clathrin (CLTA, CLTB, CLTC), AP2, PICALM, EPN2) clustering in distinct phases (early, middle and late) following LH treatment (**Figure 2C-E** and **Figure S2G**).

G_α_s interaction was highest at the earliest time point (1 min) following LH treatment, but then fell below the basal level without LH stimulation at time points beyond 3 min, a finding supported by prior BRET-based studies showing that G_α_s resides in close proximity, or “pre-coupled”, to LHR ^44,45^ (**Figure 2C-E**). Additional G_α_s-cAMP signaling machinery enriched following agonist treatment included three adenylate cyclases ADCY3, ADCY6 and ADCY9 with the latter exhibiting a transient interaction profile, and the catalytic subunits of protein kinase A (PRKACA and PRKACB) all showing up in the early interaction cluster (**Figure 2C-E**).

In contrast, LH-dependent association of LHR with β-arrestin-2 continued to increase following LH treatment and during G_α_s uncoupling, and persisted through later time points consistent with previous resonance energy transfer studies demonstrating sustained LH-dependent interactions with β-arrestin-2 ^43,44^ (**Figure 2C-E**). Upon β-arrestin-2 recruitment, LHR is known to internalize via clathrin-coated pits (CCP), and indeed interaction with clathrin heavy and light chains (CLTA, CLTB, CLTC) peaked at 2 min following LH treatment, consistent with CCP formation coinciding with β-arrestin-2 recruitment (**Figure 2C-E**). Other enriched CCP components included AP2 complexes, PICALM, EPS15 and EPN1/2, consistent with previous studies using β2AR-APEX2 ^30,31^, with EPS15/EPN1 in the early/middle and EPN2/β-arrestin-2 in the late interaction clusters sharing a similar temporal profile (**Figure 2C,D**). ADP-ribosylation factor 6 (ARF6) exhibited a peak interaction profile following 1-2 min of LH stimulation, consistent with its role regulating early events in LHR internalization upstream of the scission of clathrin-coated pits ^46^ (**Figure 2C,D**). In contrast, interaction with ARF GTPase-activating protein GIT1 was sustained after peaking at 2 min, as previously described for β2AR ^30^ (**Figure 2C,D**).

As noted above, LHR is known to traffic through VEEs within 3 min to activate acute G protein signaling and enable subsequent rapid receptor recycling ^19^. APPL1, the only adaptor protein previously shown to localize to VEEs ^19^, was not detected in these studies, likely due to limitations of abundance and sensitivity of mass spectrometry-based proteomics. However, the PDZ domain protein GIPC1 ^42^, which is necessary for LHR sorting into VEE ^20^, was enriched in agonist-treated samples, with a temporal profile strongly aligned with prior TIRF-microscopy imaging studies demonstrating GIPC1 recruitment during receptor clustering into CCPs, and dissociation following clathrin-coated vesicle disassembly and entry into VEEs ^20^ (**Figure 2C-E**).

### Leveraging the LHR-GIPC1 temporal profile to reveal novel LHR-APEX2 interactions

To demonstrate the potential for APEX2 proximity proteomics to identify new prospective regulatory interactors in a currently poorly understood pathway, we focused on previously unknown LHR-APEX2 interactors at the early endocytosis transition to VEEs. Given the requirement for GIPC1 to dissociate from LHR prior to VEE entry ^20^, the peak of LHR-GIPC1 interaction was used as a reference point for the start of VEE entry (**Figure 2F**). We applied a four-step bioinformatics data analysis workflow to identify potential new VEE-associated proteins with a peak enrichment right-shifted to later time points than GIPC1 (**Figure 3**). After initial filtering, normalization and significance analysis steps described before in **Figure 2A** (scheme repeated in **STEP 1** (**Figure 3**)), triplicate enrichment data for 2439 proteins were first visualized with principle component analysis (PCA), volcano plots and hierarchical clustering (**Figure 3A**). PCA demonstrated the high correlation between replicate time points and clear differences between early time points (1, 2 min) and later time points (5, 10 min) per each replicate (**Figure S5A**). Volcano plots highlighted the differences in enrichment of relevant selected proteins over the “no LH” control (“0”) for each of the time points of LH treatment (refer to **Figure S5B-F** and **Data S2** for quantitative data), demonstrating that each of these proteins was significantly enriched (p =0.05) at least in one of the time points. Hierarchical clustering of the cross-replicate (**Figure S6A**) and averaged (**Figure S6B**) normalized protein abundance distributed proteins into five clusters with distinct patterns of enrichment (refer to **Data S3** for quantitative data). Clusters A and B showed proteins with transient/early LHR interaction patterns with proximity peaking at 1-2 min; Clusters C and D followed with middle/late interactions and Cluster E was enriched with late/sustained proximity interactions (**Figure S6C-G**). Hierarchical clustering was insufficient to clearly identify hits in our study due to the data complexity and very subtle changes between time-points, therefore the next analysis steps were developed. In **STEP 2**, 2439 proteins were distributed into two groups arranged by the difference between their abundance values at the 2- and 3-min time points as shown on a difference plot (**Figure 3B** and **Figure S5G**). Proteins in the first group peaked at the 1- or 2-min time points, and proteins in the second group peaked at or after the 3 min time point. In **STEP 3**, data for the 577 proteins in the second group were extracted and subjected to categorical filtering by GO term “ENDO”, performed in Perseus ^39^, resulting in 191 proteins bearing this term (**Figure 3C**). The most enriched categories after analysis were “clathrin-coated vesicle”, “coated membrane”, and “vesicle coat”. To define a focused subset of hits for functional studies, these 191 proteins were subjected to manual selection based on their likely functional relationship to VEEs in **STEP 4**. These included proteins with association to core endosomal functions such as endosome fusion, acidification and membrane protein sorting (e.g. Rabs, sorting nexins, V-type ATPases), cAMP generation and cAMP/PKA effectors or role in signalosomes (e.g. adenylate cyclases, AKAPs, Ras-related GTPases), and association with cytoskeletal machinery and peripheral vesicles (e.g. ARF6 GTPase activating proteins). This led to 45 selected hits arranged by their interaction clusters with enriched VEE-related cellular component function indicated (**Figure 3D** and **Figure S6H**), which were taken forward for further study.

**Figure 3.**
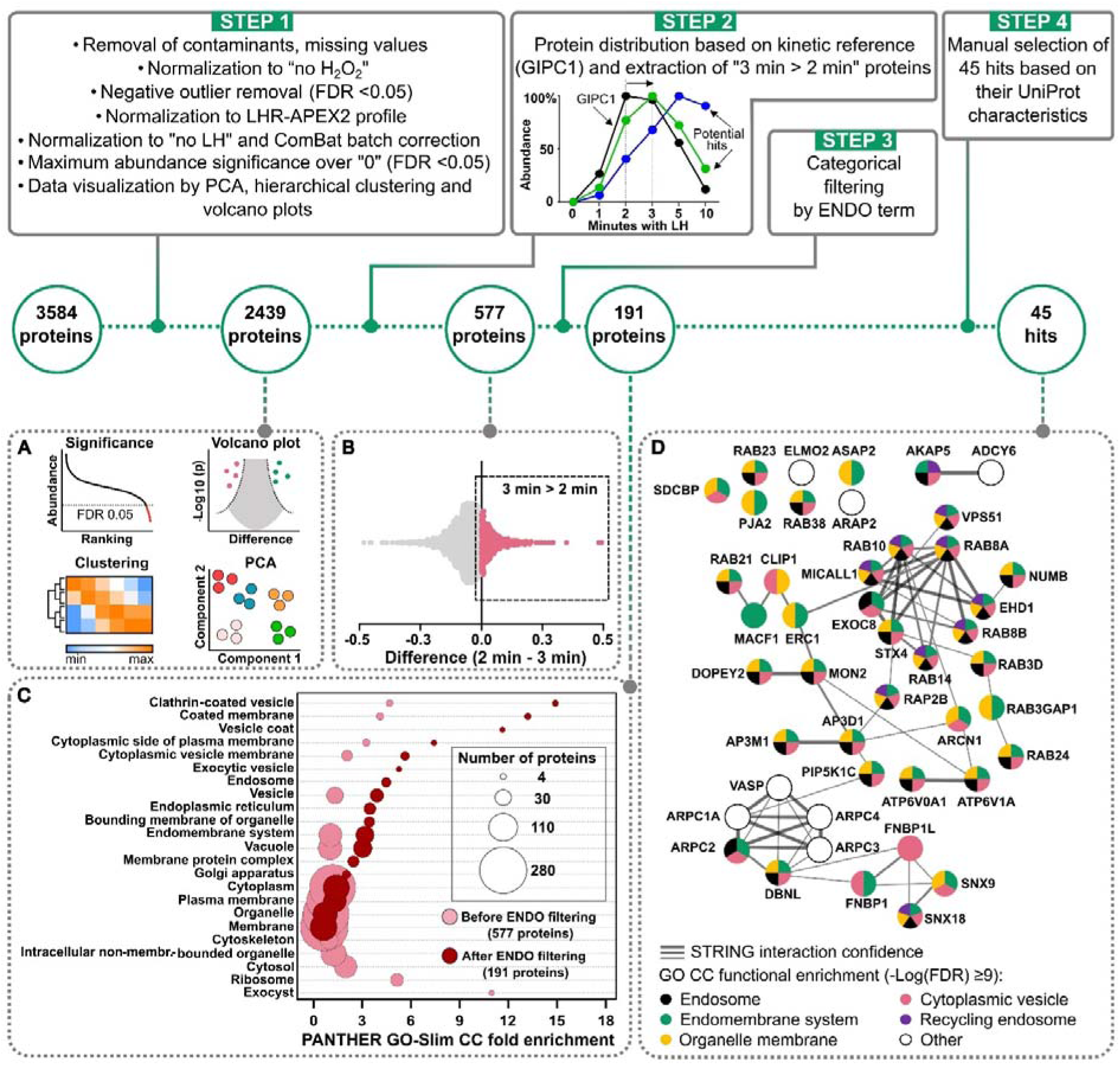
Interactomics data analysis workflow based on the GIPC1 temporal reference profile. **STEP 1.** For the details of protein filtration and normalization steps refer to Figure 2A legend. **A.** Data for 2439 proteins were visualized with PCA, volcano plots and hierarchical clustering (Refer **to Figures S5, S6** and **Data S2, S3**). **STEP 2**. Using the GIPC1 temporal profile as a reference (inset graph), data for 2439 proteins were distributed in two groups arranged by the difference between their abundance values at the 2 and 3 min time points, as shown on the difference plot (**B**), and 577 proteins in the “3 min > 2 min” group were extracted. **STEP 3**. 577 proteins were subjected to categorical filtering by term “ENDO” across GOCC, BP, MF and KEGG categories, performed in Perseus, resulting in 191 proteins bearing this term (**C**). The graph shows PANTHER GO-Slim cell component category enrichment value for proteins before (577, pink) and after (191, burgundy) filtering step; the bubble size corresponds to the number of proteins. **STEP 4**. After filtering, 191 proteins were subjected to manual selection based on their likely functional relationship to VEEs, leading to 45 selected hits. Panel **D** shows 45 selected hits arranged by their interaction clusters derived from STRING web-based platform ^47^ (figure adapted from Cytoscape (3.10.0) ^48^). The edges represent STRING database score. The enriched GO CC (Cellular Component) function for each protein (derived from STRING; (-Log(FDR) ≥9)) likely related to VEE is indicated with different coloured slices. A more detailed review of the data analysis workflow is provided in the Supplementary Materials and Methods. For the full list of proteins originated from each analysis step (**A**-**D**), with their enrichment levels at different time points, refer to the **Data S1**.

### Identification of functional interactors of LHR modulating G_α_s signal transduction

To functionally annotate potential modulators of LHR signaling we screened all 45 selected hits and 5 controls (non-targeting control (NTC), β-arrestin-2 (ARRB2), the VEE proteins APPL1 and GIPC1 and G_α_s (the essential mediator of G_α_s-coupled signaling, GNAS)) by RNA interference (RNAi) knock-down for changes in cAMP produced by WT LHR, under LH stimulation. FLAG-LHR WT HEK293 stable cells were reverse transfected with an siRNA library and grown for 96 hours. Screens were performed at two different LH concentrations to maximize the detection window for knock-downs that increase cAMP levels (3 nM LH, Screen 1, **Figure 4A**) or reduce cAMP levels (10 nM LH, Screen 2, **Figure 4B**). To focus our analysis on knock-downs which did not cause major disruption in cellular function, each well was additionally analyzed manually by phase contrast microscopy, and instances with altered cell viability or morphology were not considered further (shown as discarded points on **Figure 4A-B**). GNAS knock-down was used as a positive control for signal inhibition and exhibited nearly 100% reduction in LH-mediated cAMP generation, as expected for the G_α_s-coupled LHR. The highest-ranking hits found in both screens which reduced cAMP generation on knock-down, and therefore are potential *positive* modulators of LHR signaling, were RAP2B, EHD1, ARPC1A, MON2 with additional ARPC3, ELMO2, DBNL, SNX18, AP3M1 found in either Screen 1 or 2; the highest-ranking hits that increased cAMP generation on knock-down in both screens were RAB38, RAB23, ATP6V0A1, SNX9 with additional AKAP5, AP3D1, RAB24, MACF1 found in either Screen 1 or 2, and were considered potential *negative* modulators of signaling.

**Figure 4.**
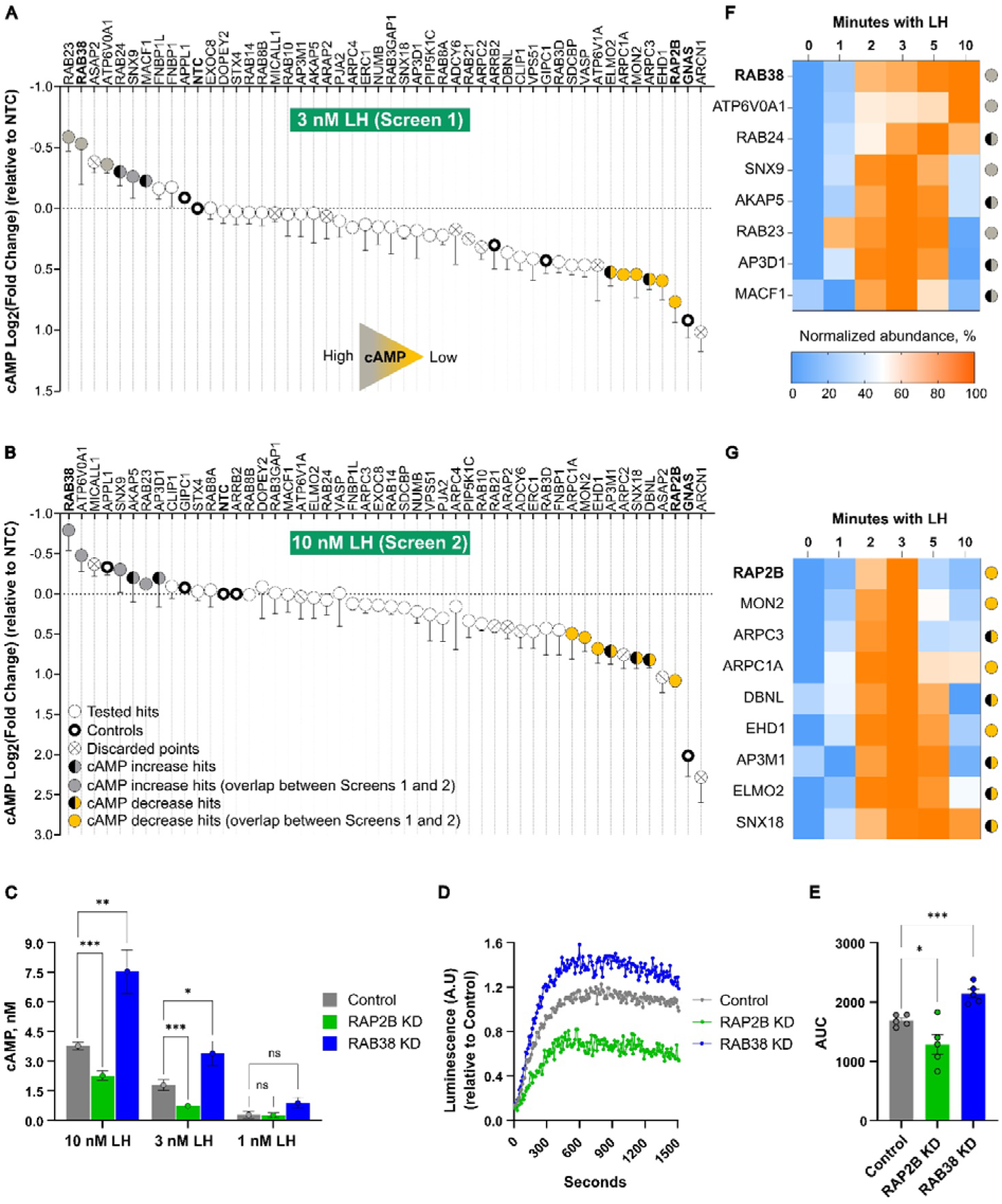
Identification of functional interactors of LHR modulating G_α_s signal transduction. FLAG-LHR HEK293 stable cells were transfected with an siRNA library for 96 h and stimulated with LH for 5 min (3 nM LH for Screen 1 (**A**) and 10 nM LH for Screen 2 (**B**)). Intracellular cAMP levels were measured from cell lysates by a TR-FRET cAMP assay. Controls included a non-targeting control (NTC) siRNA, and siRNA for GNAS (the essential mediator of G_α_s-coupled signaling), the VEE proteins APPL1 and GIPC1, and β-arrestin-2 (ARRB2). Responses were normalized to the NTC value per each replicate, transformed to Log2 and the plot is ranked from high cAMP (grey) to low cAMP (yellow). Hits found in both Screens 1 and 2 are shown as grey and yellow circles; hits found in either Screen 1 or 2 are shown as grey/black and yellow/black circles. Hits which exhibited negative effects on cell morphology and/or viability after knock-down are shown as discarded points (cross). The most changed hits present in both screens were RAP2B and RAB38. Data represent mean ± SEM (n = 3). **C**. cAMP generated by RAP2B and RAB38 knock-downs (KD) compared to NTC (Control) upon stimulation with 1 nM, 3 nM or 10 nM LH for 5 min. cAMP concentration was interpolated from TR-FRET signal using a cAMP standard curve. Data represent mean ± SEM (n = 3). Statistical analysis was performed by unpaired Student’s t-test: ns > 0.05, *p < 0.05, **p < 0.01, ***p < 0.001. See also Figure **S7A** for cAMP dose-response curves. **D,E**. Live cAMP production indicated by GloSensor ^49^ luminescence over 30 minutes (shown for over 25 minutes) by RAP2B and RAB38 knock-downs (KD) following stimulation with 10 nM LH. Representative kinetic trace, values normalized to NTC (Control) measurement at 30 minutes (**D**). Data represent mean of the area under the curve (AUC) ± SEM (n = 5). Statistical analysis was performed by unpaired Student’s t-test: *p < 0.05, ***p < 0.001. (**E**). **F**. Heatmap of interaction profiles for knock-downs leading to increased cAMP (hits are putative *negative* regulators of LHR signaling); protein enrichment across timepoints was first normalized to unstimulated (“no LH”) control, followed by normalization to the maximum and minimum enrichment signal for each protein. The profiles show a trend of a sustained interaction with LHR, with interactors staying in proximity after 3 and 5 min. **G**. Heatmap of interaction profiles for knock-downs leading to decreased cAMP (hits are putative *positive* regulators of LHR signaling). Profiles show a trend of a sharp interaction with LHR between 2-3 min upon LH stimulation.

We next further validated the impact on LHR-mediated cAMP production for knock-down of the two strongest validated cAMP-modulating hits, RAP2B and RAB38, following stimulation with 1 nM, 3 nM or 10 nM LH (**Figure 4C**). The LH-dependent cAMP levels produced by RAP2B knock-down cells was ∼2.5 times less than in control cells, whereas for RAB38 knock-down the increase was ∼2.4 times compared to the control cells, suggesting a significant effect on agonist-mediated LHR signaling function. To further assess LHR signaling, we measured cAMP dose-response curves under two knock-down conditions, that showed significant increase (∼150% for RAB38) and decrease (∼35% for RAP2B) in efficacy of LH-medicated cAMP production compared to Control (100%) (**Figure S7A**). The impact on the kinetics of cAMP signaling was assessed by employing the cAMP GloSensor^TM^ reporter ^49^ which demonstrated that the increase (via RAB38 knock-down) or decrease (via RAP2B knock-down) in agonist-dependent LHR cAMP signaling was sustained over an extended period post-stimulation with LH (**Figure 4D,E**). To rule out possible effects of knock-downs on LHR surface expression, which could affect the cAMP response, we performed flow cytometry, that showed no effect on LHR expression for either knock-down (**Figure S7B**). Comparison of heatmaps of interaction profiles for knock-downs leading to cAMP decrease or increase indicated that LHR-APEX2 interactors with a potential positive effect on LHR signaling (i.e. which lead to decreased cAMP signaling on knock-down) exhibit a profile with a peak of association at 3 min followed by dissociation, whereas the majority of interactors with a potential negative regulatory effect (i.e. which lead to increased cAMP signaling on knock-down) exhibit sustained proximity to LHR up to and beyond 5 min post-stimulation with LH (**Figures 4F,G**).

### RAP2B knock-down reroutes LHR to early endosomes

RAP2B is a member of the Ras-related superfamily of small GTPases, with a known role as a *downstream* effector of cAMP/EPAC signalosomes ^50^. Activation of RAP2B by EPAC leads to phospholipase C-epsilon signaling, but RAP2B has not previously been reported to function upstream of the cAMP pathway in generating, promoting or maintaining cAMP signaling. We have previously demonstrated that acute LHR-cAMP signaling requires receptor internalization ^19^, and to further clarify a potential novel role of RAP2B we explored the impact of RAP2B knock-down on LHR internalization and trafficking by confocal microscopy. FLAG-LHR HEK293 stable cells were transfected with non-targeting control (NTC), GIPC1 or RAP2B siRNA and plated onto PDL-coated coverslips 72 h post-transfection. After a further 24 h, cells were treated with 10 nM LH for 5 min, stripped, fixed, and labeled with secondary anti-FLAG and anti-EEA1 antibodies, followed by confocal microscopy imaging. It is known that following 10 min LH treatment, LHR rapidly internalizes primarily to a VEE compartment, and that disruption of this sorting via loss of GIPC1 redirects receptors to the early endosome and to a degradative pathway ^20,42^. Therefore, colocalization of LHR with the early endosomal marker EEA1 was measured following knock-down of RAP2B or GIPC1 to assess the impact on LHR endosomal trafficking. Interestingly, knock-down of either RAP2B or GIPC1 increased colocalization of LHR with EEA1 by approximately 2-fold compared to NTC (Control) (**Figure 5A,B**). These data suggest that in the absence of RAP2B LHR is rerouted to early endosomes, implying that RAP2B may play a role in promoting LHR trafficking to the VEE (**Figure 5C**).

**Figure 5.**
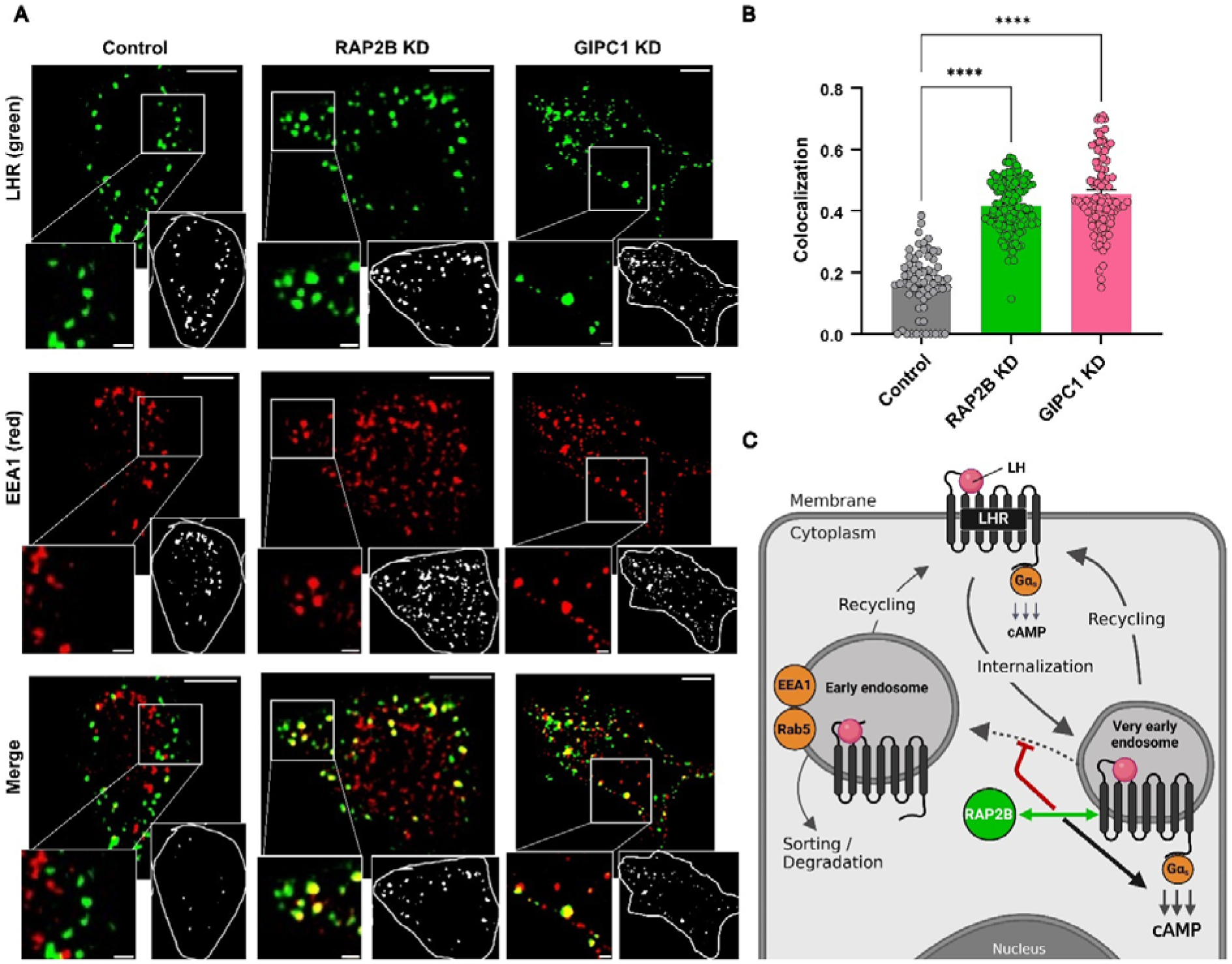
RAP2B knock-down reroutes LHR to early endosomes. **A**. Confocal images of FLAG-LHR HEK293 stable cells under three knock-down (KD) conditions: non-targeting control, RAP2B and GIPC1. FLAG-LHR was labeled with anti-FLAG antibody and treated with 10 nM LH for 10 min. Cells were stripped, fixed, permeabilized and stained with an anti-EEA1 antibody, followed by secondary AF594- and AF488-conjugated antibodies. Shown is a representative frame from imaging of cells via confocal microscopy with a zoom-in inset (bottom left). A representative single cell output image with threshold mask for each channel and colocalized pixels is shown in another inset (bottom right). Scale bar = 5 µm. Scale bar in the zoom-in inset = 1 µm. **B**. Quantification of colocalization between LHR channel (green) and EEA1 channel (red) that translates to how many LHR-containing endosomes are also EEA1-positive. The Manders colocalization coefficient is shown under three knock-down conditions. Z-slices were considered separately; n = 10 cells per condition; 15 z-slices per each cell. Data represent mean ± SEM. Statistical analysis was performed by unpaired Student’s t-test: ****p < 0.0001. **C**. A mechanistic hypothesis for the role of RAP2B interaction with LHR in very early endosomes (VEE). Our data suggest that RAP2B may have direct roles in stimulating LHR cAMP signaling from VEE as well as inhibiting fusion of LHR-containing VEE with early endosomes.

## Discussion

Quantitative proximity proteomics ^28,29^ has recently emerged as a cutting edge technology to capture the complexity of GPCR networks at a spatial and temporal level ^30–32,36,51^. However, interrogation of these activation-dependent interactomes resolved on a scale of minutes can be challenging, raising significant demands on robust data analysis ^29^, and to date this approach has been mostly applied to well-understood GPCRs with a largely established interactome. In this study, we applied quantitative APEX2 ^22^ proximity proteomics to define the LHR interaction network on a minute-to-minute timescale. We developed an integrated experimental and analytical methodology which combines highly reproducible quantitative TMT-based proteomics with temporal reference profiles for key interactors to identify novel LHR interactors in a specific spatial compartment, the VEE ^19,20^. Although applied here to LHR, the pipeline developed could be applicable to any GPCR which undergoes rapid early and post-endocytic events, and/or where a receptor traverses multiple subcellular compartments.

Using LH stimulation time points based on established LHR signaling kinetics, endocytosis and trafficking to and from VEEs ^19,20^, proximity proteomics identified almost ∼2500 LH-dependent interactions, which is a similar range to previously reported studies applying APEX2 ^30,35^. Only a subset of these proteins is likely to be direct interactors of LHR with the potential to functionally regulate LHR activity, since APEX2 labeling also captures indirect interactors within a 20 nm radius as well as ‘bystander’ proteins. A robust analytical pipeline was essential to overcome several confounding features of proximity labeling. First, the relative abundance of a particular protein in the interactome does not necessarily strongly correlate with physical interaction; for example, GIPC1 and clathrin heavy chain were similarly enriched (**Figure S2F**) despite the former having a known direct interaction with LHR ^20,42^. Second, bona fide interactors may not be identified due to the intrinsic limitations of mass spectrometry to detect low abundance proteins or which lack proteotypic tryptic peptides. For example, APPL1 was not identified in this study. Third, the shortest temporal resolution of APEX2 is currently limited to ca. 1 min, and therefore may not fully capture events at even shorter timescales. Fourth, as a membrane protein, LHR-APEX2 would be expected to be present in various subcellular compartments (e.g. ER, Golgi) in addition to the plasma membrane and endosomes, potentially contributing to the challenges of interactome deconvolution ^52^. Fifth, given the high labeling promiscuity, APEX2 also captures ‘bystanders’ freely diffusing through the proximity cloud, usually leading to the identification of thousands of captured proteins. This caveat can be circumvented by the use of spatial references, however, this approach was not feasible in this study due to the lack of well-defined organelle locations for LHR. Finally, affinity purification also generates background due to nonspecific protein interactions with the resin. Therefore, TMT-labeled controls (“no H_2_O_2_” and “no LH”) and a cascade of filtering and normalization steps were required to provide a cut-off and select high confidence LHR-interacting proteins for further analysis (**Figure 2A**). All protein profiles were normalized to the LHR-APEX2 abundance profile as an internal standard per replicate, to account for any variations during labeling and sample processing, which proved to be the critical step in our analysis as demonstrated by the representative protein profiles before and after normalization (**Figure S2B-E**).

Our refined pipeline approach robustly captured both direct and indirect known LHR interactors with excellent spatiotemporal resolution, including several with transient or low-affinity interactions, such as G protein signal transducers (**Figure 2** and **Figure S3**), β-arrestin-2 and ARF6 (**Figure 2**). Interestingly, members of all four heterotrimeric G protein families were identified in the screen (**Figure S3**). LHR is a Gαs (GNAS)-coupled GPCR, with a G protein signal pathway attributed to many of its physiological functions. Prior studies employing resonance energy transfer approaches have also demonstrated pre-association of LHR with GNAS as well as rapid and transient ligand-mediated interactions ^44,45^. As for many GPCRs, the LHR is also known to exhibit pleiotropic signaling properties, including activation of Gαq/11 and Gαi/o signaling ^53–55^. Our study corroborated the previous findings in these coupling profiles, further supporting the rigor of the interactomic profile identified for this GPCR. Although a potential link of LHR to cytoskeleton remodeling has been suggested through its direct interaction with ARNO/ARF6 complex and activation of RAC1 ^53^, we are not aware of prior studies reporting a potential association/coupling of LHR with Gα12/13 family members. In addition, this dataset has provided further information on specific subtypes that could associate with each of these G protein families with potential distinct ligand-induced temporal profiles.

The HEK293 cell model also enabled identification of interactors specifically relevant to physiological LHR signaling, including Hippo signaling components involved in ovarian granulosa cell to luteal cell differentiation ^56^ (**Figure S4**). We focused on functional validation of VEE compartment interactions in the present work due to the importance of the VEE for LHR signaling and trafficking. However, the dynamic interactomic datasets generated here may also prove a valuable resource for future mapping of the landscape of LH-induced LHR interactions at the plasma membrane, during formation of CCPs, and other rapid activation-dependent events which are currently poorly characterized for this receptor.

The application of a temporal profile for a known interactor (GIPC1 in the present study) as an internal reference for functional annotation of dynamic networks when GPCR pharmacological tools (i.e. agonists and antagonists) are limited, offers a powerful alternative to other approaches which employed comparative proteomics (e.g., agonist vs antagonist; agonist vs biased agonist ^30,36^) or spatial references at specific locations (e.g. plasma membrane, endosome, etc) to discriminate interactors form ‘bystander’ proteins ^31,57–59^. Spatial references represent a powerful approach to overcome the issue of ‘bystanders’, however, they require complex analysis across distantly related APEX2 datasets ^31^ and are not feasible when the protein of interest lacks well-defined organelle location at a given time. In fact, for LHR the endosomal properties and cellular machinery of the VEE is poorly understood, with the additional challenge that these endosomal networks are interconnected ^20^, thus prompting the distinct analytical approach described in this study. Previous biochemical and cell biology studies have demonstrated that GIPC1 interaction with the LHR C-terminal tail occurs during LHR clathrin-mediated endocytosis, dissociating prior to entering the VEE compartment ^20^ (**Figure 2C**). The exquisite temporal resolution of APEX2 labeling enabled us to exclude LHR interactors with maximum abundances at timepoints prior to GIPC1 dissociation and focus on plausible regulators of VEE trafficking and/or signaling (**Figure 3**). Interestingly, our functional studies identified two contrasting interaction profiles, with putative positive regulators of LHR signaling showing interaction profiles peaking at 3 min post-activation (**Figure 4G**), whereas the majority of putative negative regulators maintained their proximity from 3 min across the 10 min ligand stimulation window of the experiment (**Figure 4F**). This temporal profile of negative regulators may suggest such LHR complexes function to switch off signal activity prior to receptor recycling back to the plasma membrane, given LHR recycling has been captured within this same time frame ^19^.

Two novel putative LHR interactors, RAP2B and RAB38 were found as positive and negative regulators of LHR signaling, respectively, with RAP2B also regulating endosomal distribution of LHR. RAB38 is a member of the family of Rab small GTPases, well known for their roles in regulating membrane trafficking, and has been previously studied in the context of melanosome biogenesis ^60,61^. Prior studies of the classic early endosomal RAB5 demonstrated no role in VEE generation or impact on LHR trafficking ^20^, and the identification of a distinct Rab family member in this study which may promote LHR-mediated cAMP signaling from the VEE is a significant step towards a mechanistic understanding of VEE-dependent signaling, and an important topic for future studies. RAP2B is a member of the Ras-related family of small G proteins. While this protein has been reported to be a downstream regulator of cAMP signaling from certain G_α_s-coupled GPCRs, where the cAMP effector EPAC with RAP2B activates calcium signaling via PLC-epsilon ^62^, this is the first report of a potential role for RAP2B upstream of cAMP generation. In addition, RAP2B exhibits parallels with the known activity of GIPC1 in retaining LHR within the VEE. However, in contrast to GIPC1, RAP2B also seems to be a positive regulator of LHR-cAMP signaling, a novel function which may be independent from any potential ability to regulate endosomal sorting of LHR. Future studies could assess the alteration in these complex networks following knockdown of known (GIPC1) or putative interactors (RAP2B, RAB38) of LHR, akin to recent proximity proteomic studies with the 5HT2A serotonin receptor ^57^.

Overall, we present an APEX2 pipeline optimized for the study of GPCRs, and likely also other membrane receptors, with minimal existing pharmacological tools, spatial markers and/or known protein interactors. Our findings highlight the intricate complexity of GPCR interactomes and illustrate the importance of developing methods to deconvolute this complexity. We expect that these and similar approaches will enable broader applicability of proximity labeling to accelerate fundamental discovery research for the large superfamily of GPCRs.

### Resource availability

The mass spectrometry proteomics data have been deposited to the ProteomeXchange Consortium via the PRIDE partner repository with the dataset identifier PXD053246. Reviewer access details: Log in to the PRIDE website using the following details: Project accession: PXD053246; Token: XLs7p7S5LAIs. Alternatively, reviewer can access the dataset by logging in to the PRIDE website using the following account details: Username; Password: zmLRrH3u9kaz

## Acknowledgements

We acknowledge funding support from the Laboratory for Synthetic Chemistry and Chemical Biology under the Health@InnoHK Program of The Government of Hong Kong Special Administrative Region of the People’s Republic of China and the Biotechnology and Biological Sciences Research Council (BBSRC), grants BB/S001565/1 and BB/V006142/1. Schematics were created in BioRender.com. We would like to thank Prof. Mark von Zastrow (Univ. California, San Francisco, USA), Prof Gunther Schmalzing (RWTH Aachen Univ., Germany), Prof Ali Tavassoli (Univ. Southampton, UK) and Dr Kim Jonas (King’s College London, UK) for providing certain plasmids. We would like to thank Shabbir Ahmed (Leica Microsystems) and Tracey Williams (Leica Microsystems) for access and assistance with Mica (Zeiss) imaging. We would like to thank Tate group members for assistance and insightful discussions and Erika Bernardini for help with cell culture. We would like to acknowledge Dr Roman Fedoryshchak for his assistance with confocal microscopy and critical reading of this manuscript.

## Author contributions

MMS, EWT and ACH conceptualized the study, obtained funding and provided resources for the study. RR contributed to confocal microscopy studies. ARW carried out GloSensor assay and analysis. MAS carried out flow cytometry and measured dose-response curves under knock-down conditions. MMS developed and carried out all other experiments in this study, developed bioinformatics analysis pipeline, performed functional studies, confocal microscopy, data analysis and made all the figures. DC supervised proteomics set up and provided guidance. JH contributed to some of the proteomics analysis steps and generated the Cytoscape figure panel. MMS, ACH and EWT wrote the original draft of the manuscript, and all authors provided comments, reviewed, and edited the final manuscript.

## EXPERIMENTAL MODEL AND STUDY PARTICIPANT DETAILS

### Cell culture

Flp-In T-Rex 293 cell line was obtained from Francis Crick Institute, UK and used for all proximity-labeling experiments. Prior to stable transfection, cells were cultured in high-glucose, no pyruvate Dulbecco’ Modified Medium (DMEM) + GlutaMAX (Gibco), supplemented with 10% heat-inactivated fetal bovine serum (FBS, Gibco) and 100 μg/mL Zeocin (Invitrogen). Upon stable transfection, Zeocin was removed, and cell medium was supplemented with 5 μg/mL Blasticidin (Thermo Scientific). For selection, 100 μg/mL Hygromycin B (Invitrogen) was added in addition to Blasticidin and all stable cells were kept under these antibiotics. All cells were maintained in a humidified incubator at 37°C and 5% CO_2_.

## METHOD DETAILS

### Generation of pcDNA5-FRT-TO-LHR-APEX2 plasmid

The LHR-APEX2 construct was assembled in the following order: a preprolactin sequence, a FLAG tag, a human LHR sequence without a stop codon, a short 4 amino acid linker and an APEX2 sequence with a stop codon at the end. The whole construct was inserted into a pcDNA5-FRT-TO vector containing two tetracycline repressor sites upstream of the construct (**Figure S1A**).

The APEX2 gene was linearized by PCR from the pcDNA3.1 plasmid (kind gift from Dr Braden Lobingier, USA) using primers cgagtctagaGGAAAGTCTTACCCAACTGTGAG (forward, overlaps with a reverse LHR primer) and tcccggtgtcttctatggagttaGGCATCAGCAAACCCAAG (reverse, overlaps with a forward vector primer) with a stop codon included in the end.

The hLHRwt cDNA together with an upstream preprolactin signal sequence (ATGGACAGCAAAGGTTCGTCGCAGAAAGGGTCCCGCCTGCTCCTGCTGCTGGTGGTGTCAAATCTACTCTTGTGCCAGGGTGTGGTCTCC), followed by a FLAG tag was amplified by PCR from the pcDNA3.1 plasmid (as described in ^20^) using primers ttagtgaaccgtcagatcgcATGGACAGCAAAGGTTCG and aagactttccTCTAGACTCGAGACACTCTG. The pcDNA5-FRT-TO vector (kind gift from Prof Gunther Schmalzing, Germany) was linearized by PCR using primers CTCCATAGAAGACACCGGGAC (forward, overlaps with a reverse APEX2 primer) and GCGATCTGACGGTTCACTAAAC (reverse, overlaps with forward LHR primer). The pcDNA5-FRT-TO-LHR-APEX2 fusion sequence was assembled with NEBuilder HIFI DNA Assembly Master Mix (New England Biolabs).

### Generation of stable Flp-In T-Rex 293 LHR-APEX2 cells

Flp-In T-Rex 293 cells were seeded onto 10 cm dishes and grown to 75% confluency. Cells were then transfected with a mixture of two plasmids, pOG44 (kind gift from Ali Tavassoli, UK) and pcDNA5-FRT-TO-LHR-APEX2 at a ratio of 9:1 (8 μg DNA total) using 48 μL of Lipofectamine 2000 (Invitrogen) in Opti-MEM (Gibco) following the manufacturer’s protocol. After 24 h post-transfection, media was changed to a fresh one. After 48h post-transfection, cells were split and grown to 40-50% confluency. Next, media was replaced with complete DMEM supplemented with 100 μg/mL Hygromycin B (Invitrogen) to start the selection of stable clones. In 10 days, selection was complete.

### Generation of stable FLAG-LHR WT HEK 293 cells

The generation of this cell line has been previously described in ^20^.

### Proximity labeling

Flp-In T-Rex 293 LHR-APEX2 cells were passaged a few times after thawing. For all replicates, cells were of the same passage. For each replicate, cells were plated into 7 separate 10 cm dishes per each labeling condition and treated with 1 μg/mL of tetracycline hydrochloride (Merck) overnight to induce LHR-APEX2 protein expression. Each treatment apart from H_2_O_2_ and quenching was done in a humidified incubator at 37°C and 5% CO_2_. *It should be noted here, that all treatments were set in a way that no more than three plates ended up being H_2_O_2_-treated and then quenched at the same time due to a complexity of the handling*.

Media was removed and all cells were treated with 5 mL of 500 μM biotin phenol (Tocris Bioscience) in complete media for exactly 30 min before quenching. Next, cells were treated with LHR agonist LH (50 nM final concentration) to initiate LHR signaling. For that, 1 mL of LH (300 nM) was added directly on top of plates and cells were incubated for 1, 2, 3, 5, and 10 min before H_2_O_2_ addition. Two control plates were treated with no LH media for 1 min. Within 15 min of addition, 2 mM H_2_O_2_ solution in ice-cold PBS was prepared and stored on ice. 6 mL of 2 mM H_2_O_2_ solution was added on top of each plate (apart from one control plate) at the 29^th^ minute of BP treatment, and the plates were gently rotated. A countdown timer was set right at the H_2_O_2_ addition. After 55s, media in each plate was gently but quickly poured away into a beaker. After 60s, all plates were quenched with 4 mL of an ice-cold quenching buffer containing 10 mM sodium azide (Merck), 10 mM sodium ascorbate (Merck) and 5 mM Trolox (Merck) in DPBS, repeated 3 times. Cells were then scraped off in a quenching buffer, centrifuged and pellets collected. Pellets could be stored at -80°C for a few weeks.

### Protein precipitation

All buffers were prepared fresh before each experiment and filtered through 0.22 μm filters. The Eppendorf Protein LoBind microcentrifuge tubes were used for all steps. All solvents were proteomics grade. Cell pellets were thawed on ice. 100 μL of supplemented RIPA buffer (10 mM sodium azide, 10 mM sodium ascorbate, 5 mM Trolox, 1mM PMSF, 1x cOmplete Protease Inhibitor (Roche) in RIPA, pH 7.5) was added to each pellet, samples were resuspended and left on ice for 20 min. Lysates were clarified in a tabletop centrifuge (13000 rpm, 10 min, 4°C). Supernatants were collected and *could be flash-frozen at this point to be stored at -80°C for a few days.* Proteins were precipitated by addition of methanol (200 μL), chloroform (50 μL) and water (100 μL) to the 100 μL of each lysate, followed by centrifugation in a microcentrifuge (6000 rpm, 4 min, RT). The upper layer was aspirated, followed by addition of methanol (200 μL), vortexing, sonication and centrifugation (13000 rpm, 3 min, RT). The resulting protein pellet was again x2 washed with methanol (200 μL), vortexed, sonicated, centrifuged and air-dried for 2-3 min. *Samples could be frozen at this stage (-20°C) for overnight before decanting of final methanol wash solution.* Protein pellets were resuspended in 100 μL of 1% SDS (w/v) in HEPES (50 mM, pH 8) and sonicated until the pellets broke completely. Solutions were centrifuged (13000 rpm, 3 min, RT) to make sure no pellet formed. Each sample was diluted with 400 μL of HEPES (50 mM, pH 8) for the final solution of 0.2% SDS (w/v) in HEPES (50 mM, pH 8). Solutions were centrifuged (13000 rpm, 3 min, RT) to make sure no pellet formed. *Samples could be frozen at -20°C at this point for overnight*. Protein concentrations were measured using the BCA assay.

### Western blotting of biotinylated proteins

10 μg of protein per sample dissolved in 0.2% SDS (w/v) in HEPES (50 mM, pH 8) was used. Samples were mixed with 2-mercaptoethanol-supplemented Laemmli Sample buffer (Bio-Rad), boiled for 10 min and separated by 12% SDS-PAGE gel. Gels were transferred to a nitrocellulose membrane (Bio-Rad) and blocked with 3% (wt/vol) BSA in Tris-buffered saline with 0.1% Tween-20 (TBS-T) for 2h at RT. Next, membranes were incubated with neutravidin-HRP conjugate (Thermo Fisher Scientific) in blocking buffer (1:2000) for 1h at RT. Membranes were washed with TBS-T (x4, 5 min). The bands of biotinylated proteins were detected by western fluorescent detection reagent (Millipore Luminata Western HRP Chemiluminescence Substrate) and images were recorded with the ImageQuant LAS 4000 series.

### Streptavidin pull-down, reduction, alkylation and digestion of biotinylated proteins

All buffers were prepared fresh before each experiment and filtered through 0.22 μm filters. The Eppendorf Protein LoBind microcentrifuge tubes were used for all steps. All solvents were proteomics grade. For the pull-down, 600 μg of protein per sample was used. The corresponding amount of sample was transferred to a new tube, topped up to 500 μL with 0.2% SDS/HEPES (50 mM, pH 8) and applied to 60 μL of agarose neutravidin beads slurry (Thermo Fisher Scientific) (beads were previously decanted of the storage buffer, x3 washed with 0.2% SDS/HEPES and redissolved in 0.2% SDS/HEPES). The samples were incubated for 4h at RT in a shaker. Next, the tubes were centrifuged (13000 rpm, 4 min, RT), the beads collected, x3 washed with 900 μL of 0.2% SDS/HEPES and washed three times with 900 μL of HEPES. The supernatants after the first centrifugation were later used to assess the pull-down efficiency by western blotting. The beads in each sample were resuspended in 150 μL HEPES (50 mM, pH 8), whereas 5 μL of slurry per sample were set aside to assess pull-down efficiency by western blotting. Next, 15 μL of reducing agent tris(2-carboxyethyl)phosphine hydrochloride (TCEP) (Merck) (5 mM final) and 15 μL of alkylating agent chloroacetamide (CAA) (Merck) (15 mM final) were added and samples were incubated for 10 min at RT in a shaker. 0.5 μg of LysC (Promega; 20 μg vial dissolved in 105 μL of HEPES (50 mM, pH 8)) and 1 μg of Sequencing Grade Modified Trypsin (Promega; 20 μg vial dissolved in 105 μL of HEPES (50 mM, pH 8)) were added to each sample, then gently vortexed, briefly centrifuged and incubated in a thermoshaker (1000 rpm, 18h, 37°C).

### Pull-down efficiency analysis by western blotting

5 μL of bead slurry per sample, 5 μL of supernatant and 10 μg of protein sample before pull-down dissolved in 0.2% SDS (w/v) in HEPES (50 mM, pH 8) were mixed with 2-mercaptoethanol-supplemented Laemmli Sample buffer (Bio-Rad), boiled for 10 min and separated by 12% SDS-PAGE gel. All next steps were performed as described above for the **western blotting of biotinylated proteins.**

### The Tandem Mass Tag (TMT) labeling

The Eppendorf Protein LoBind microcentrifuge tubes were used for all steps. All solvents were proteomics grade. The beads were briefly centrifuged and the reaction in each sample was quenched with 6 μL of 50x EDTA-free cOmplete protease inhibitor cocktail (Roche), gently vortexed and centrifuged again. The supernatants were transferred to fresh tubes, the beads were washed with 160 μL of HEPES and two supernatant fractions combined. 320 μL of each sample were transferred to a new tube and the liquid was evaporated on a SpeedVac for 4h at 45°C. The leftover 10-20 μL per sample were used to measure peptide concentration by Pierce Quantitative assay (Thermo Fisher Scientific) following the manufacturer’s protocol, which was in the range of 80-100 μg/mL. The dried samples were resuspended in 200 μL of proteomics grade water and the pH was checked to be 8. Only half (100 μL) of each sample (which equals to ∼ 15 μg of peptides) was used for labeling, the rest was dried on a SpeedVac for 4h at 45°C and frozen at -80°C. TMT10plex reagents set (Thermo Fisher Scientific; 0.8 mg per vial) were thawed and each reagent dissolved in 105 μL of dry acetonitrile. Each TMT reagent was aliquoted at 20 μL (∼160 μg, sufficient amount to label 25μg of peptides per sample) into 3 separate tubes (according to the number of replicates). Identical samples across replicates were labeled with the reagents of the same mass tag according to the table below. For that, 100 μL of peptide samples were added on top of TMT reagents according to the table below and the reactions were incubated for 1h at RT in a shaker.

|  |  |  |  |  |  |  |  |  |
| --- | --- | --- | --- | --- | --- | --- | --- | --- |
| No H<sub>2</sub>O<sub>2</sub> | No LH | 1 min | 2 min | 3 min | 5 min | 10 min |  | **Replicate 1** |
| 126 | 127N | 127C | 128C | 129C | 130C | 131 |  |  |
| No H<sub>2</sub>O<sub>2</sub> | No LH | 1 min | 2 min | 3 min | 5 min | 10 min |  | **Replicate 2** |
| 126 | 127N | 127C | 128C | 129C | 130C | 131 |  |  |
| No H<sub>2</sub>O<sub>2</sub> | No LH | 1 min | 2 min | 3 min | 5 min | 10 min |  | **Replicate 3** |
| 126 | 127N | 127C | 128C | 129C | 130C | 131 |  |  |

The reactions were quenched by addition of 1.7 μL of 5% hydroxylamine and left to incubate for 15 min at RT in a shaker. All samples <u>per replicate</u> were then combined in one tube and the liquid was evaporated on a SpeedVac for 18h at RT.

### Sample fractionation

The samples were redissolved in 300 μL of 0.1%TFA / water and acidified to pH ∼3 by addition of a few μL of TFA. The fractionation of samples was done with a Pierce High pH Reversed-Phased Kit (Thermo Fisher Scientific) following the manufacturer’s protocol. Briefly, the sample solutions were sonicated, vortexed and briefly centrifuged. Each sample was loaded onto a pre-conditioned cartridge, centrifuged in a microcentrifuge (6000 rpm, 3 min, RT) and the eluent collected. In the same manner, cartridges were subsequently washed with a series of freshly prepared eluting solutions with gradually increasing concentrations of acetonitrile. This resulted in eight separate fractions in total per each sample that were evaporated on a SpeedVac for 18h at 35°C. The dried peptides could be stored at -80°C for a few weeks.

### nLC-MS/MS analysis

The peptides were dissolved in 10 μL of resuspension solution (Optima LC/MS grade water (Fisher Scientific) containing 2% (v/v) UPLC grade acetonitrile and 0.5% (v/v) TFA), shaken (5 min, RT), sonicated (5 min, RT), vortexed and centrifuged (13000 rpm, 3 min, RT). Samples were filtered through 3x Durapore membrane filters (Millipore) plugged into a p20 pipette tip by centrifuging the samples through the filters (4000 × g, 5 min) into a mass spectrometry vial. Samples were stored at 4 °C until ready for analysis. 3 μL of each filtered sample were injected and separated on an EASY-Spray Acclain PepMap C18 column (EASY-nLC 1000) (50 cm x 75 μm inner diameter, Thermo Fisher Scientific) using a 3h gradient separation of 0–100% solvent B (80% MeCN supplemented with 0.1% formic acid): solvent A (2% MeCN supplemented with 0.1% formic acid) at a flow rate of 250 nL/min. EASY-nLC 1000 was coupled to a QExactive mass spectrometer via an easy-spray source (Thermo Fisher Scientific). The QExactive was operated in data-dependent mode with survey scans acquired at a resolution of 70,000 at m/z 200 (transient time 256 ms). Scans were acquired from 350 to 1800 *m/z*. Up to 10 of the most abundant isotope patterns with charge +2 or higher from the survey scan were selected with an isolation window of 1.6 m/z and fragmented by higher-energy collision dissociation (HCD) with normalized collision energies of 25 W. The maximum ion injection times for the survey scan and the MS/MS scans (acquired with a resolution of 35,000 at m/z 200) were 20 and 120 ms, respectively. The ion target value for MS was set to 10^6^ and for MS/MS to 10^5^, and the intensity threshold was set to 8.3 × 10^2^.

### Data analysis

Peptide searches were performed in MaxQuant (version 1.6.10.43) ^38^. Under group-specific parameters and type, reporter ion MS2 was selected, and the appropriate TMT10plex isobaric labels selected for both lysines and N-termini. The isotope errors contained in each TMT batch code was also entered. For all experiments, oxidation (M) and acetyl (protein N-term) were set as variable modifications, carbamidomethyl (C) was set as a fixed modification, trypsin/P was set as the digestion mode. The N-terminal acetylation delta masses were 42.01056; the TMT-10plex delta masses were 229.16293. Where multiple TMT sets were analysed, re-quantify and match between runs were selected. The latest UniProt FASTA files for the human proteome and contaminants databases were used.

Data analysis was performed in Perseus ^39^ versions 1.5.0.9 and 1.6.1.2. Reporter intensity corrected values were loaded into the matrix. Data was filtered by removing rows based on “reverse”, and “potential contaminant” columns. Proteins with total TMT intensity values >100 were removed. Data was log2 transformed, missing values (NaN) were removed retaining those rows that had all valid values at least in two replicates out of three. Missing data (6.4%) were imputed under a missing-at-random assumption using a knn method ^63^.

Data was normalized to “no H_2_O_2_” control (background) and protein rows with log2 fold enrichment values significantly enriched over background were identified in a Significance A outliers test (FDR <0.05 cut-off) in Perseus (**Figure 2A**). Data was internally normalized to LHR-APEX2 profile across three replicates to account for variance in protein abundance across samples (**Figure S2B-E**). For that, “subtract row cluster” function in Perseus was used. Data was then normalized to the no agonist control (“no LH”) and corrected for inter-plex batch variation using ComBat ^64^. Maximum log2 fold enrichment values across all time points per replicate were subjected to a t-test (Benjamini-Hochberg FDR <0.05) in Perseus to identify significantly enriched proteins following agonist treatment. The full list of proteins with quantitative data can be found in **Data S1** table.

For functional enrichment of Kyoto Encyclopedia of Genes and Genomes (KEGG) pathways in the dataset, DAVID 2021 ^41^ (https://david.ncifcrf.gov/) web-based software was employed (**Figure 2B**). Quantification of enriched pathways was performed in GraphPad Prism (GraphPad Software).

The proteins were distributed into two groups by comparing their maximum abundances at “2 min” and “3 min” time points in Perseus and visualized on a difference plot in GraphPad Prism (**Figure 3B**). The protein pathways and categories were annotated by the KEGG and Gene Ontology (GO) (GO biological process (GOBP), GO molecular function (GOMF) and GO cellular component (GOCC)) in Perseus, followed by categorical/pathway filtering based on the “ENDO” search word. To demonstrate this step visually, the lists of proteins before and after enrichment were uploaded to Gene Ontology web-based database and GO statistically significant enrichment analysis performed using the PANTHER Overrepresentation test (PANTHER version 18.0) and PANTHER GO-Slim Cellular Component against a reference list containing all Homo sapiens genes in the database (Fisher’s Exact test, FDR p <0.5 cut-off). The GO-Slim Cellular Component categories were plotted in GraphPad Prism based on their fold enrichment values alongside the number of proteins in these categories (**Figure 3C**). 191 proteins were subjected to manual selection based on their likely functional relationship to VEEs as described in the main text.

To visualize protein hits network clustering, the list was uploaded to STRING ^47^ web-based database ((https://string-db.org/)) and networks were visualized using Cytoscape (version 3.10.0) ^48^ (https://cytoscape.org) with edges representing STRING database score. Colour slices represent the enriched GO Cellular Component functions derived from STRING ((-Log(FDR) ≥9)) (**Figure 3D**).

Comparison of log2 protein enrichment at various time points with agonist to no agonist control was performed in Perseus and visualized by volcano plots. After initial filtering and normalization steps described above, the log2 protein enrichment values were mean averaged and subjected to a Student’s t-test (p <0.05) to identify significantly enriched proteins (**Figure S5** and **Data S2**).

Hierarchical clustering was performed in Perseus using the built-in clustering algorithm, the optimal number of clusters was identified by the Elbow method. After initial filtering and normalization steps described above, the log2 protein enrichment values were normalized to their maximum and minimum enrichment signal (scale 0 to 1), clustered and visualized using Microsoft Excel (**Figure S6** and **Data S3**).

Unless mentioned otherwise, all quantification and visualization of data was done in GraphPad Prism and Excel.

### cAMP signaling assay

Intracellular cAMP was determined by homogenous time-resolved fluorescence (HTRF) LANCE *Ultra* cAMP assay (Perkin Elmer). For the EC50 value measurements, Flp-In T-Rex 293 LHR-APEX2 cells induced overnight with 1 μg/mL of tetracycline hydrochloride (Merck), non-induced cells or FLAG-LHR WT HEK293 cells grown in dishes were detached with trypsin, centrifuged and pellets redissolved in complete cell medium (Flp-In T-Rex 293 LHR-APEX2 cells: high-glucose, no pyruvate DMEM + GlutaMAX supplemented with 10% FBS and antibiotics as described above; FLAG-LHR WT HEK293 cells: high-glucose, no pyruvate DMEM (Sigma Aldrich) supplemented with 10% FBS). Cells were counted (in six replicates and average cell number used) and diluted to 200 cells/μL in FBS-free cell medium right before the assay. Cells were transferred to a white OptiPlate-384 microplate (Perkin Elmer) at 5 μL per well in triplicates per each ligand concentration. Next, various concentrations of LH dissolved in FBS-free cell medium were added at 5 μL per each well. For unstimulated condition, just 5 μL of FBS-free cell medium was added. Cells were stimulated for 5 min at 37°C, followed by addition of 5 μL of Eu-cAMP tracer and 5 μL of ULight-anti-cAMP antibody dissolved in lysis buffer according to the manufacturer’s protocol. The plate was covered and incubated for 1h at RT in the dark and read on an EnVision Multimode Plate reader (Perkin Elmer) equipped with a UV2 320 nm excitation filter and two emission filters (203 Eu 615 nm and 205 APC 665 nm).

For the EC50 value measurements under Control, RAP2B and RAB38 knock-down conditions, the HTRF-based cAMP Gαs dynamic kit (Revvity) was employed. The FLAG-LHR WT HEK293 stable cells plated in 96 well plates were stimulated with LH as above and reaction was terminated by aspirating the media and placing the plates on ice followed by addition of 60 μL ice-cold lysis buffer. 10 μL cell lysates were transferred to a white OptiPlate-384 microplate. 5 μL of the cAMP Eu-cryptate-labeled antibody and 5 μL of the cAMP-d2 reagent were added to each well according to the manufacturer’s protocol. The plate was covered and incubated for 1h at RT in the dark and read on a PHERAstar FSX plate reader (BMG Labtech Ltd) equipped with a 337 nm excitation filter and 620/665 nm dual emission filters. Data was normalized to protein concentration per sample quantified by the Bradford Assay (Themo Fisher Scientific). All data points were normalized to the LHR WT E_max_ under Control knock-down conditions at Log –7 (considered 100% maximum response) per each experiment. All quantification analysis was performed using GraphPad Prism (GraphPad Software). All error bars represent SEM based on at least 3 biologically independent experiments.

### siRNA reverse transfection

An ON-TARGETplus siRNA SMARTpool Cherry-pick library of 54 targets (49 hits and 5 controls) in a 96-well plate (0.25 nmole siRNA per well) was purchased from Horizon Discovery/Dharmacon (Cambridge). The siRNA library plate was resuspended in 50 μL of 1X siRNA buffer (Horizon Discovery) to make 5 μM stock and aliquoted. The FLAG-LHR WT HEK293 stable cells were plated at 10,000 cells per well into a 96-well siRNA library plate and reverse transfected with 100 nM siRNA using DharmaFECT 1 transfection reagent (Horizon Discovery) in Opti-MEM (Gibco) following the manufacturer’s protocol. Briefly, 2 μL of 5 μM siRNA per well was mixed with 8 μL of Opti-MEM, followed by addition of 0.12 μL of DharmaFECT 1 dissolved in 10 μL of Opti-MEM. The mixture was incubated for 20 min at RT, followed by addition of 10,000 cells in 80 μL of complete DMEM. Cells were grown for 5 days (day 1 was the day of reverse transfection) and then cAMP screening assay was performed on day 5.

### cAMP siRNA screening assay

On day 5 after reverse siRNA transfection, cell media was aspirated from each knock-down well using a multichannel. 50 μL of Versene solution (Gibco) was added to each well and cells were incubated for 3 min at 37°C. 100 μL of complete DMEM was added per well and cells were carefully resuspended with a multichannel to break up any cell clots. Cells in the control NTC well were counted in six replicates and average cell number used. 30 μL of cell suspension per well was then transferred to another 96 well plate and diluted to 80 cells/μL in FBS-free cell medium right before the assay. Cells were transferred to OptiPlate-384 microplate (Perkin Elmer) at 5 μL per well (400 cells/well) in triplicates per each condition. Next, various concentrations of LH (3 nM for Screen 1 and 10 nM for Screen 2) dissolved in FBS-free cell medium were added at 5 μL per each well. For unstimulated condition, just 5 μL of FBS-free cell medium was added. Cells were stimulated for 5 min at 37°C, followed by addition of 5 μL of Eu-cAMP tracer and 5 μL of ULight-anti-cAMP antibody dissolved in lysis buffer according to the manufacturer’s protocol. The plate was incubated for 1h at RT and read on an EnVision Multimode Plate reader (Perkin Elmer) equipped with a UV2 320 nm excitation filter and two emission filters (203 Eu 615 nm and 205 APC 665 nm). All quantification analysis was performed using GraphPad Prism (GraphPad Software). All error bars represent SEM based on at least 3 biologically independent experiments. To calculate amount of cAMP produced (**Figure 4C**), data was interpolated from the cAMP standard curve. For this experiment, cells were manually counted per each condition and cell number adjusted to 80 cells/μL (400 cells/well). All quantification analysis was performed using Prism (GraphPad Software). All error bars represent SEM. Statistical analysis was performed by unpaired Student’s t-test: ****p < 0.0001.

### GloSensor assay

FLAG-LHR WT HEK 293 stable cells in a white opaque 96-well plate (Thermo Fisher Scientific) were reverse transfected (day 1) with siRNA as described above in the **siRNA reverse transfection.** On day 4 after siRNA transfection, cells were transfected with 100 ng GloSensor 20F plasmid (Promega) per well using Lipofectamine 2000 (Invitrogen) in Opti-MEM (Gibco) following the manufacturer’s protocol. 24 hours later cell media was aspirated and replaced with 90 μL equilibration media (88% CO_2_-independent medium (Gibco), 10% FBS (Merck) and 2% GloSensor cAMP Reagent stock solution (Promega)) before incubation for 2h at 37°C. Cells were transferred to a microplate reader PHERAstar Plus (BMG Labtech) prewarmed to 37°C and incubated for 5 min. 10 basal reads using a LUM plus optic module were taken before 10 μL of 100 nM LH diluted in equilibration medium was added to cells (10 nM final concentration). Immediately after ligand addition luminescence was measured every 10 seconds for 30 min with 2-3 technical replicates per experiment. Basal values for knock-down conditions (RAP2B and RAB38) were normalized to control (NTC) basal values and this was used to normalize the ligand response to account for any differences in cell number between conditions. All quantification analysis was performed using Prism (GraphPad Software). All error bars represent SEM. Statistical analysis was performed by unpaired Student’s t-test: ****p < 0.0001.

### Flow cytometry

Flow cytometry was used to determine the effect of RAP2B and RAB38 knock-down on LHR WT cell-surface expression. The FLAG-LHR WT HEK293 stable cells in a 12-well plate were reverse transfected with 120 nM ON-TARGETplus human RAP2B siRNA SMARTpool, ON-TARGETplus human RAB38 siRNA SMARTpool or ON-TARGETplus Non-targeting pool human siRNA (all from Horizon Discovery) per well using DharmaFECT 1 transfection reagent (Horizon Discovery) in Opti-MEM (Gibco) following the manufacturer’s protocol. Briefly, 24 μL of 5 μM of each siRNA per well was mixed with 100 μL of Opti-MEM, followed by addition of 2.88 μL of DharmaFECT 1 dissolved in 120 μL of Opti-MEM. The mixture was incubated for 20 min at RT, followed by addition of 120,000 cells in complete DMEM. Cells were grown for 5 days (day 1 was the day of reverse transfection) and then flow cytometry was performed on day 5. To measure the LHR surface expression, cells were treated live with M1 anti-FLAG antibody (Sigma Aldrich) (1:500 in FBS-free DMEM / 0.1% BSA) for 30 min at 37°C. Media was removed, cells washed with ice-cold PBS (x3), resuspended in PBS / 2% FCS, scraped off and centrifuged (1000 rpm, 10 min, 4°C). Pellets were resuspended in anti-mouse AF488 conjugated secondary antibody (Invitrogen) solution (in PBS / 2% FCS) and treated for 1h in the dark at 4°C. Cells were centrifuged (1000 rpm, 10 min, 4°C), supernatant removed, and pellets resuspended in PBS / 2% FCS. Plasma membrane fluorescence was quantified using the FACSCalibur Flow Cytometer (BD Biosciences) at 10,000 cells per condition.

### Immunofluorescence microscopy

For live cell imaging of FLAG-LHR and FLAG-LHR-APEX2, cells were treated live with M1 anti-FLAG antibody (Sigma Aldrich) (1:500 in FBS-free DMEM / 0.1% BSA) for 15 min at 37°C, followed by anti-mouse AF488 conjugated secondary antibody (Invitrogen) for 15 min and imaged live via confocal microscopy (Leica, SP5, x63). LH (10 nM) was added and cells imaged after 10 min. For analysis of receptor trafficking in fixed cells, FLAG-LHR WT HEK 293 stable cells in a 96-well plate were reverse transfected (day 1) with siRNA as described above in the **siRNA reverse transfection.** On day 4, 50 μL of trypsin was added to each well, cells were detached, resuspended in 150 μL of complete DMEM and the whole amount of cell suspension was added on top of coverslips pre-coated with poly-D-lysine (Sigma Aldrich) in a 24-well plate. Cells were grown for 30h and media was aspirated. Next, cells were treated live with a mouse M1-FLAG antibody (1:500 in FBS-free DMEM / 0.1% BSA) for 15 min at 37°C, followed by direct addition of LH (10 nM final) and incubation for 10 min at 37°C. Media was removed, cells washed with cold PBS (x2), followed by stripping with 0.04M EDTA in PBS (Ca^2+^ and Mg^2+^ free) (x3) to selectively remove FLAG antibody bound to the remaining surface LHR receptors. For analysis of receptor recycling, cells were plated on glass coverslips in a 12 well plate and ‘fed’ live with anti-FLAG M1 antibody (1:500 in FBS-free DMEM / 0.1% BSA) for 15 min at 37°C, followed by addition of LH (10 nM final) for 20 min. Cells were then washed 3 times with 0.04M EDTA in PBS (Ca^2+^/Mg^2+^ free) and incubated in FBS-free DMEM / 0.1% BSA for 1h.

For all fixed samples, cells were fixed with 4% paraformaldehyde (PFA) in PBS for 20 min at RT, washed with PBS (x4), blocked in PBS / 2% FCS for 20 min at RT and permeabilized with PBS / 2% FCS / 0.2% Triton-X for 15 min at RT. The solution was then removed, and coverslips were incubated with a primary rabbit anti-EEA1 antibody (Invitrogen) (1:500 in PBS / 2% FCS) for 2h at RT, followed by washing with PBS / 2% FCS (x3). Cells were then incubated with anti-rabbit AF488-conjugated and anti-mouse AF594-conjugated antibodies (Invitrogen), (both 1:500 in PBS / 2% FCS) for 30 min at RT in the dark, followed by washing with PBS (x3). Coverslips were mounted on microscopy slides using Mowiol (Sigma Aldrich) supplemented with 1 mg/mL DAPI (Invitrogen) and dried in the dark overnight. Coverslips were then imaged using a TCS-SP8 confocal microscope (Leica) or Mica (Leica) with a x63 1.4 numerical aperture (NA). Leica LAS AF image acquisition software was utilized. All subsequent Lif image files were analyzed in ImageJ (NIH).

### Image analysis

For measurement of colocalization, a line was drawn along the boundary of the cell to fully surround the cell. The Mander’s colocalization coefficients between two channels (LHR and EEA1) were determined with ImageJ BIOP JACoP colocalization plugin. Z-slices were considered separately. Threshold for each channel was adjusted manually using Otsu method. ImageJ JACoP threshold mask was used to visualize overlayed pixels from two channels (**Figure 5A**). All quantification analysis was performed using Prism (GraphPad Software). All error bars represent SEM. Statistical analysis was performed by unpaired Student’s t-test: ****p < 0.0001.

## QUANTIFICATION AND STATISTICAL ANALYSIS

### Statistics

The results are presented as mean ± standard error of the mean (SEM) based on at least 3 biologically independent experiments. Analysis of statistical significance was performed using Prism (v.10.4.1, GraphPad Software) by unpaired Student’s t-test: ****p < 0.0001.

## Supplemental Data Tables Titles

**Data S1.** Relates to **Figures 2** and **3**. Data table contains a full list of proteins identified in the study, including their quantified time-course enrichment levels, and lists of proteins with their time-course enrichment data identified in each of our bioinformatics analysis pipeline steps shown on **Figure 3**.

**Data S2.** Relates to **Figure S5.** Data table contains quantification results of volcano plot analyses.

**Data S3.** Relates to **Figure S6.** Data table contains heatmap quantification and full hierarchical clustering data.

