## Supplemental figures and legends for "A spatiotemporally resolved GPCR interactome reveals novel mediators of receptor agonism"

**Figure S1**

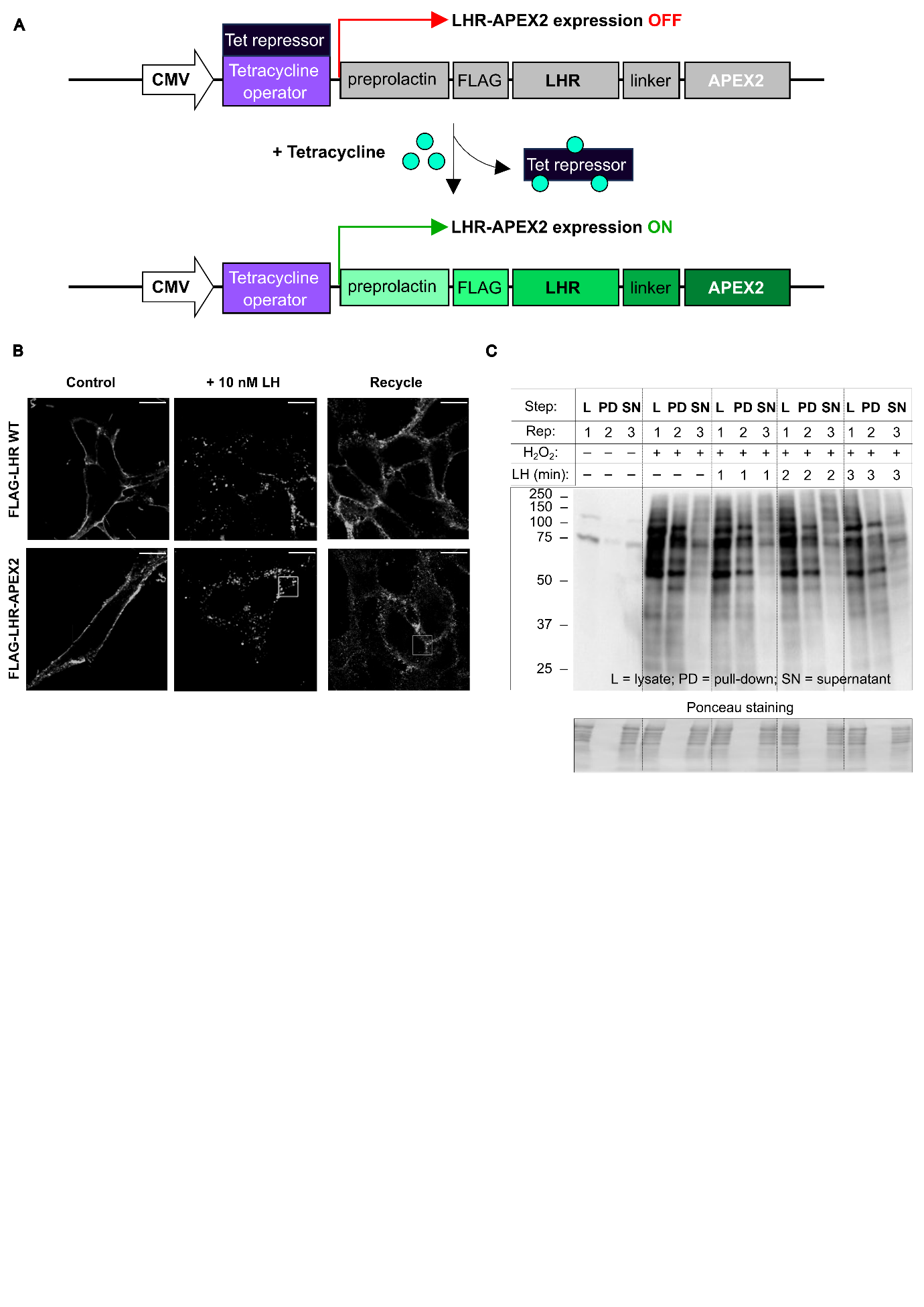

**Figure S1 (relates to Figure 1). A**. Schematics of LHR-APEX2 construct. The LHR-APEX2 construct was assembled in the following order: a preprolactin sequence, a FLAG-tag, a human wild-type LHR sequence without a stop codon, a short 4 amino acid linker and an APEX2 ^1^ sequence with a stop codon at the end. The whole construct was inserted into a pcDNA5-FRT-TO vector containing two tetracycline operator sites upstream of the construct. The construct was stably transfected into host Flp-In T-REx 293 cells ^2^. This system allows the generation of a stable cell line exhibiting tetracycline-inducible expression of a gene of interest from a specific genomic location. Addition of tetracycline “blocks” tetracycline repressor constitutively expressed in these cells, enabling expression of LHR-APEX2. **B**. Validation of LHR-APEX2 expression, trafficking, and recycling by live-cell confocal microscopy. FLAG-LHR WT HEK293 cells and FLAG-LHR-APEX2 cells were labeled with anti-FLAG antibody and non-treated (control) or treated with 10 nM LH for 10 min. Cells were then washed with PBS/0.04% EDTA followed by a further incubation for 1h to assess LHR recycling back to the plasma membrane. Cells were stained with a secondary AF488-conjugated antibody. Shown is a representative frame from imaging of live cells via confocal microscopy. Scale bar = 5 μm. Both untagged and APEX2-tagged LHR constructs exhibit comparable plasma membrane expression, LH-induced internalization into VEE and recycling. **C**. Efficiency of protein pull-down on streptavidin-coated beads accessed by neutravidin-HRP western blot. Clear changes in biotinylation levels are observed before pull-down (lysates) and after pull-down (supernatants (SN)). Pull-down (PD) samples clearly show enrichment of biotinylated proteins bands.

**Figure S2**

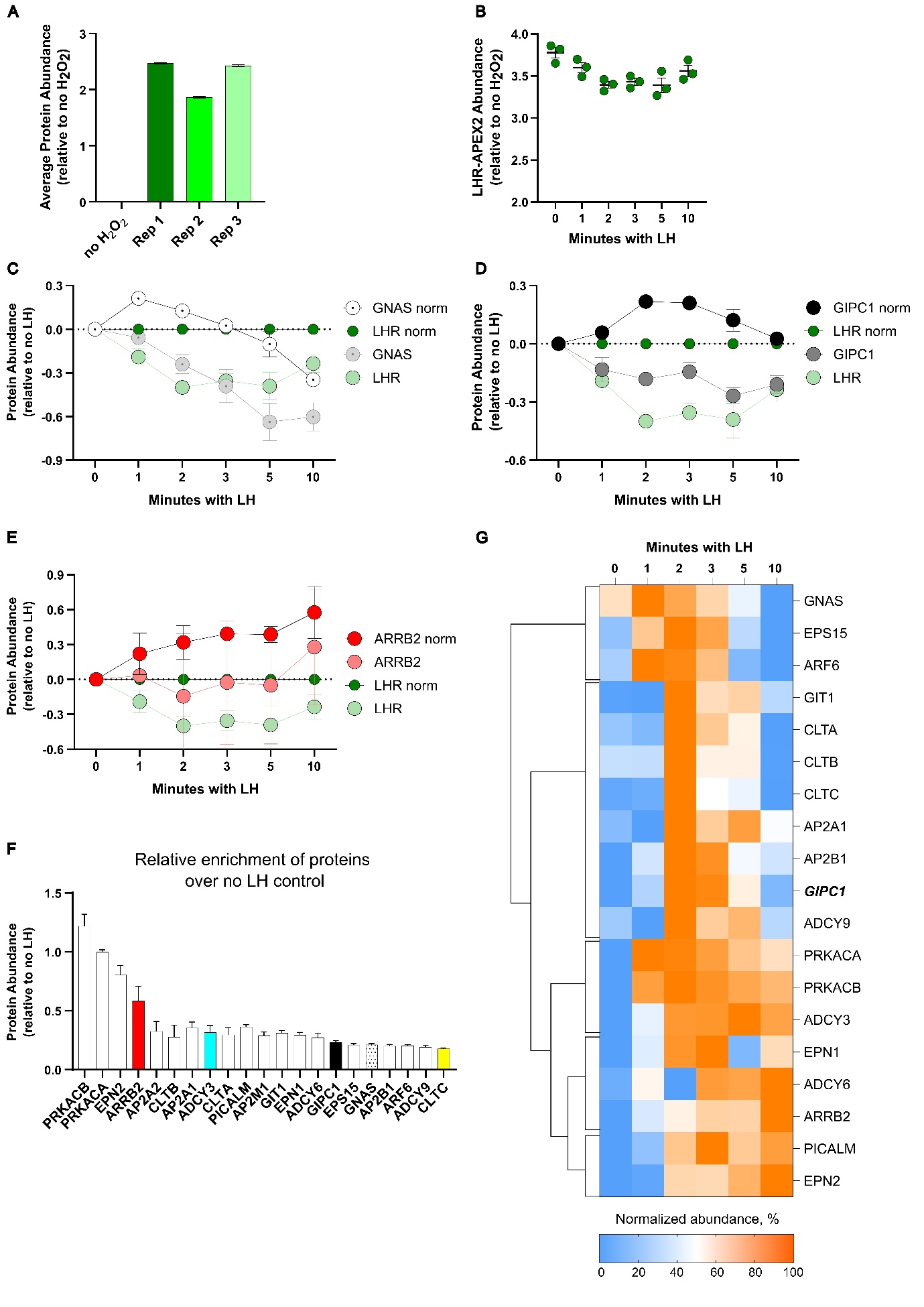

**Figure S2 (relates to Figure 2). A**. Protein abundances across three replicates normalized to “no H_2_O_2_” background control. Data shown as Log2 values (as an average of abundance values for all proteins in the same TMT experiment). **B**. Changes in LHR-APEX2 abundances normalized to “no H_2_O_2_” control in the time-course experiment with LH. Data represents mean ± SEM (n = 3). **C-E**. Representative graphs of protein temporal abundance profiles before and after normalization to the LHR-APEX2 profile that show the importance of this step for data analysis. All profiles were normalized to “no LH” control. **C**. LHR-APEX2 and GNAS profiles along time-course treatment with LH before (LHR, faded green; GNAS, faded grey) and after (LHR norm, dark green; GNAS norm, white dotted) normalization to the LHR-APEX2 profile. **D**. LHR-APEX2 and GIPC1 profiles along time-course treatment with LH before (LHR, faded green; GIPC1, faded black) and after (LHR norm, dark green; GIPC1 norm, black) normalization to the LHR-APEX2 profile. **E**. LHR-APEX2 and β-arrestin-2 (ARRB2) profiles along time-course treatment with LH before (LHR, faded green; ARRB2, faded red) and after (LHR norm, dark green; ARRB2 norm, red) normalization to the LHR-APEX2 profile. Data represents mean ± SEM (n = 3). **F**. Relative protein abundance of known LHR interactors. All protein profiles were normalized to “no LH” control and the maximum enrichment value of all time points with LH was plotted. The graph shows that abundances of direct (e.g. GIPC1) and indirect (e.g. CLTA,B,C) LHR interactors are comparable. Data represent mean ± SEM (n = 3). **G**. Heatmap of interaction profiles for known LHR interactors arranged by hierarchical clustering. Enrichment level (abundance) for each protein across different time points was normalized to the non-stimulated (“no LH”) control first, followed by normalization to its maximum and minimum enrichment signal.

**Figure S3**

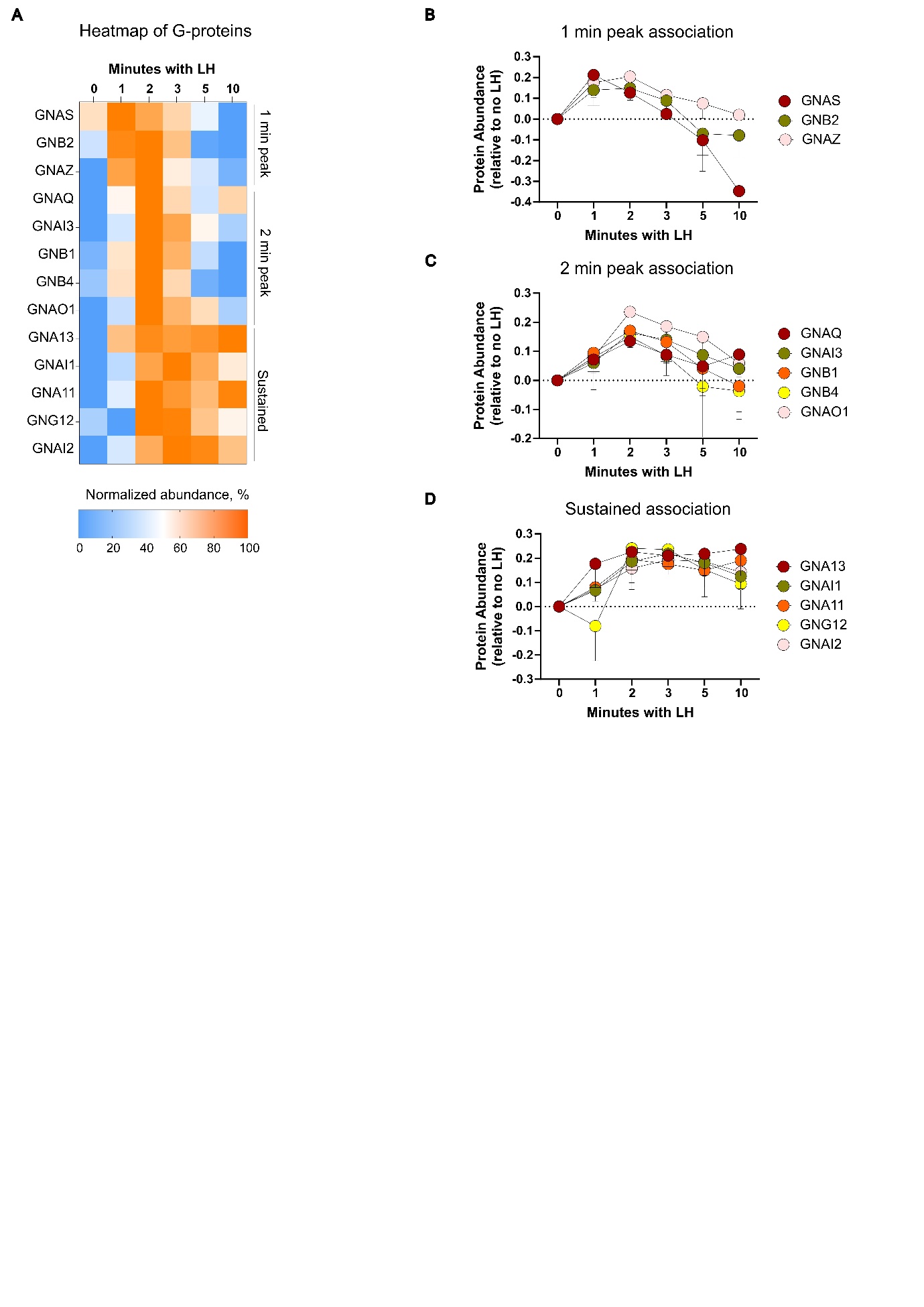

**Figure S3 (relates to Figure 2). A**. Heatmap of interaction profiles for all G proteins found in the dataset. Each protein’s enrichment level across different time points is normalized to “no LH” control first, followed by normalization to its maximum and minimum enrichment signal. The profiles are arranged by the time of their strongest interaction with LHR going from those peaking at 1 min, 2 min and to sustained G protein interactions. **B**-**D**. Temporal abundance profiles of all G proteins found in the dataset arranged by their strongest interaction with LHR such as 1 min (**B**), 2 min (**C**) and sustained association (**D**). Each protein’s enrichment level across different time points is normalized to “no LH” control. Data represents mean ± SEM (n = 3).

**Figure S4**

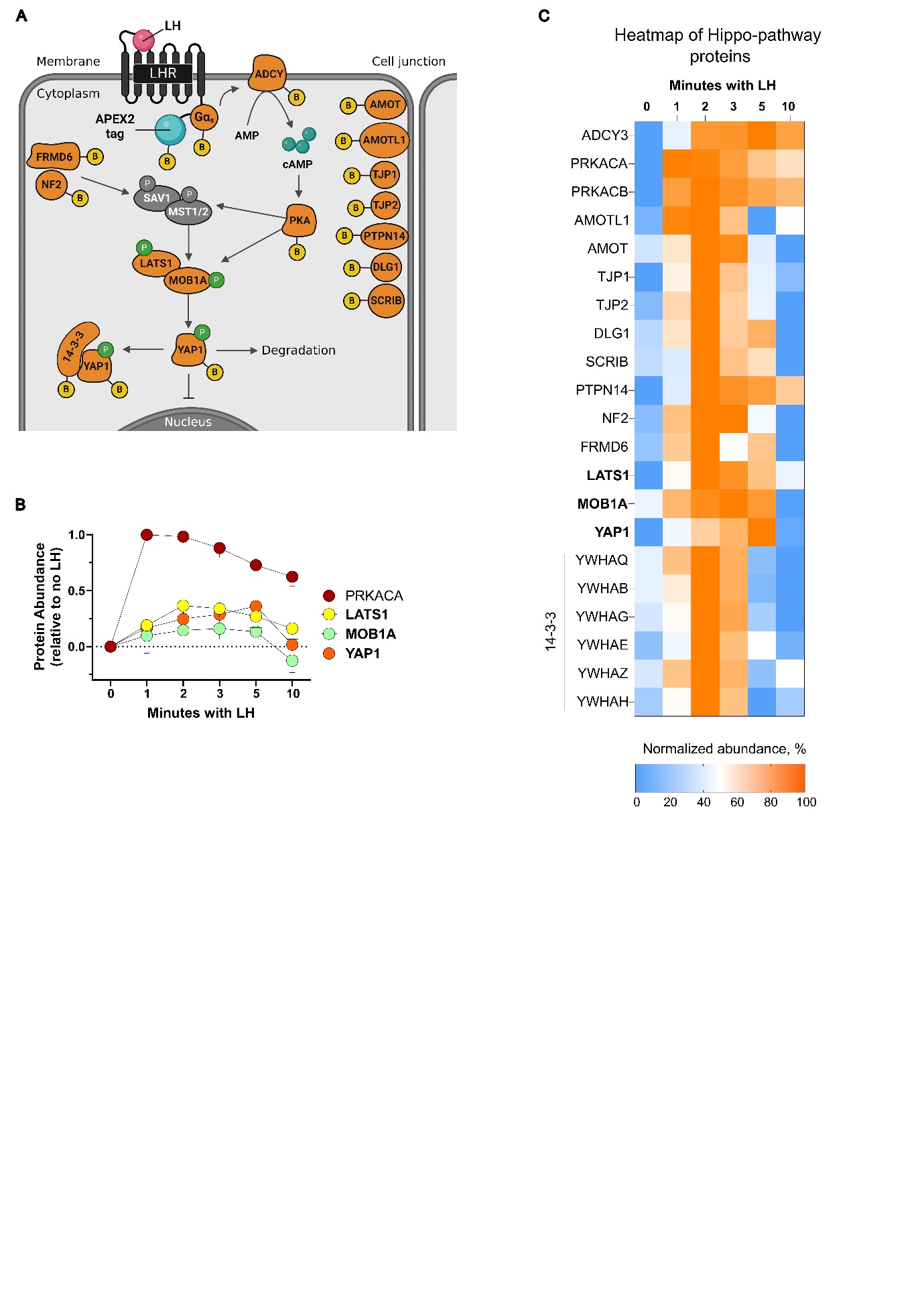

**Figure S4 (relates to Figure 2). A.** Cartoon to illustrate the Hippo-YAP1 pathway-related proteins identified by proteomics analysis. These include cell junction proteins, LATS1 and MOB1A kinases, pathway inhibitor proteins 14-3-3. SAV1 and MST1/2 kinases were not found in the dataset. **B**. Temporal profiles of selected Hippo pathway-related proteins. Each protein’s enrichment level across different time points is normalized to “no LH” control. Data represents mean ± SEM (n = 3). **C**. Heatmap of interaction profiles for the Hippo pathway-related proteins found in the dataset. Each protein’s enrichment level across different time points is normalized to “no LH” control first, followed by normalization to its maximum and minimum enrichment signal.

**Figure S5**

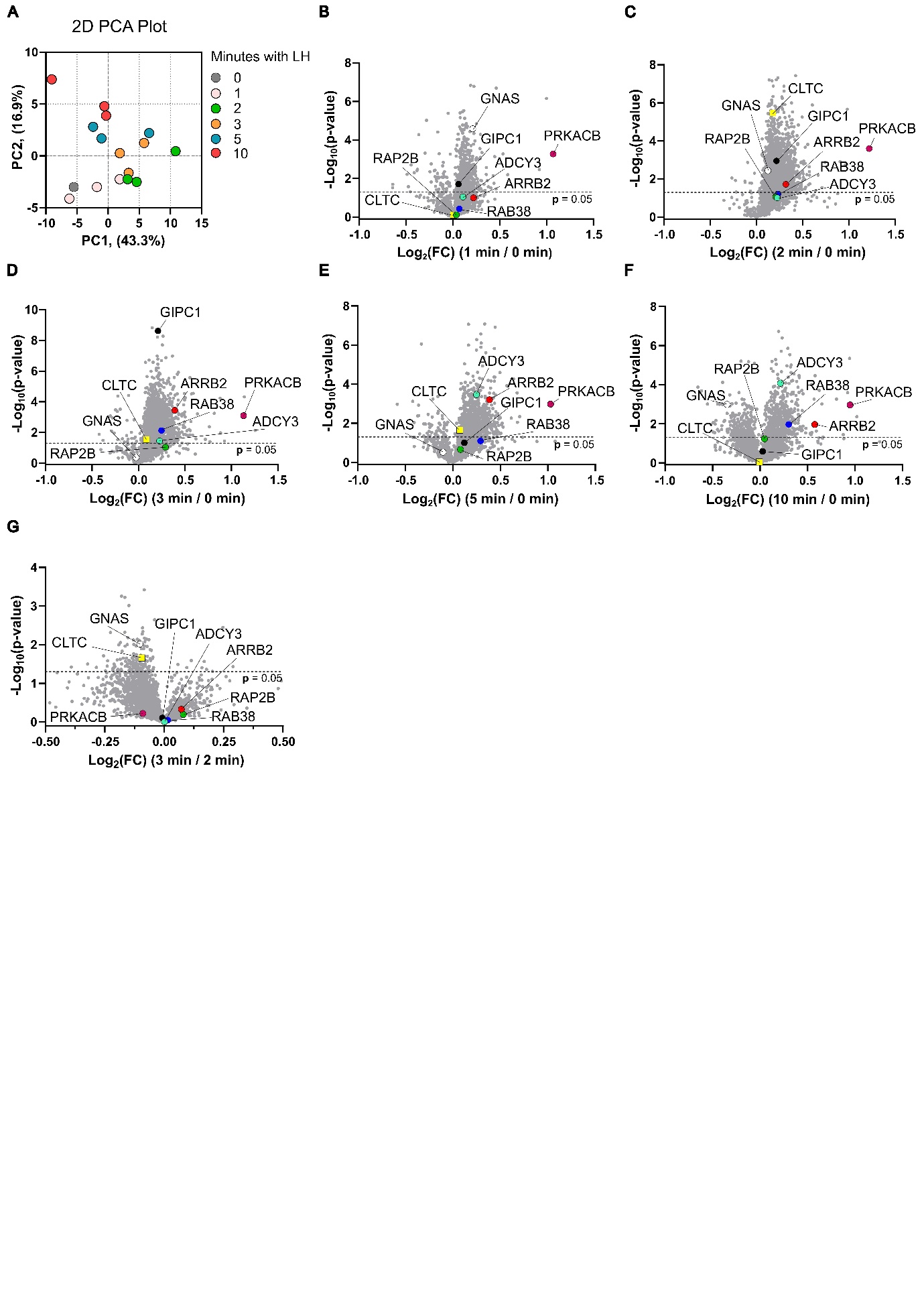

**Figure S5 (relates to Figure 3). A.** Principal component analysis (PCA) score plot shows comparison of proteomes of each replicate (n = 3) of different time points with LH. Colours indicate different LH treatment times: 0 min (grey), 1 min (pink), 2 min (green), 3 min (orange), 5 min (blue) and 10 min (red). Clustering shows high correlation between replicates and clear difference between early times points and later time points. **B**-**F**. Volcano plots comparing the enrichment of proteins at various time points with LH (1 min, **B**; 2 min, **C**; 3 min, **D**; 5 min, **E**; 10 min; **F**) vs non-stimulated samples (0 min). Selected relevant proteins are highlighted in colour: GNAS (white circle), CLTC (yellow square), ADCY3 (turquoise circle), GIPC1 (black circle), PRKACB (burgundy circle), ARRB2 (red circle), RAB38 (blue circle), RAP2B (green circle). Data represents mean (n = 3). Statistical analysis was performed in Perseus ^3^. **G**. Volcano plot comparing the enrichment of selected proteins between 2 min and 3 min treatment with LH. Data represents mean (n = 3). Statistical analysis was performed in Perseus. See also **Data S2** for volcano plot data tables.

**Figure S6**

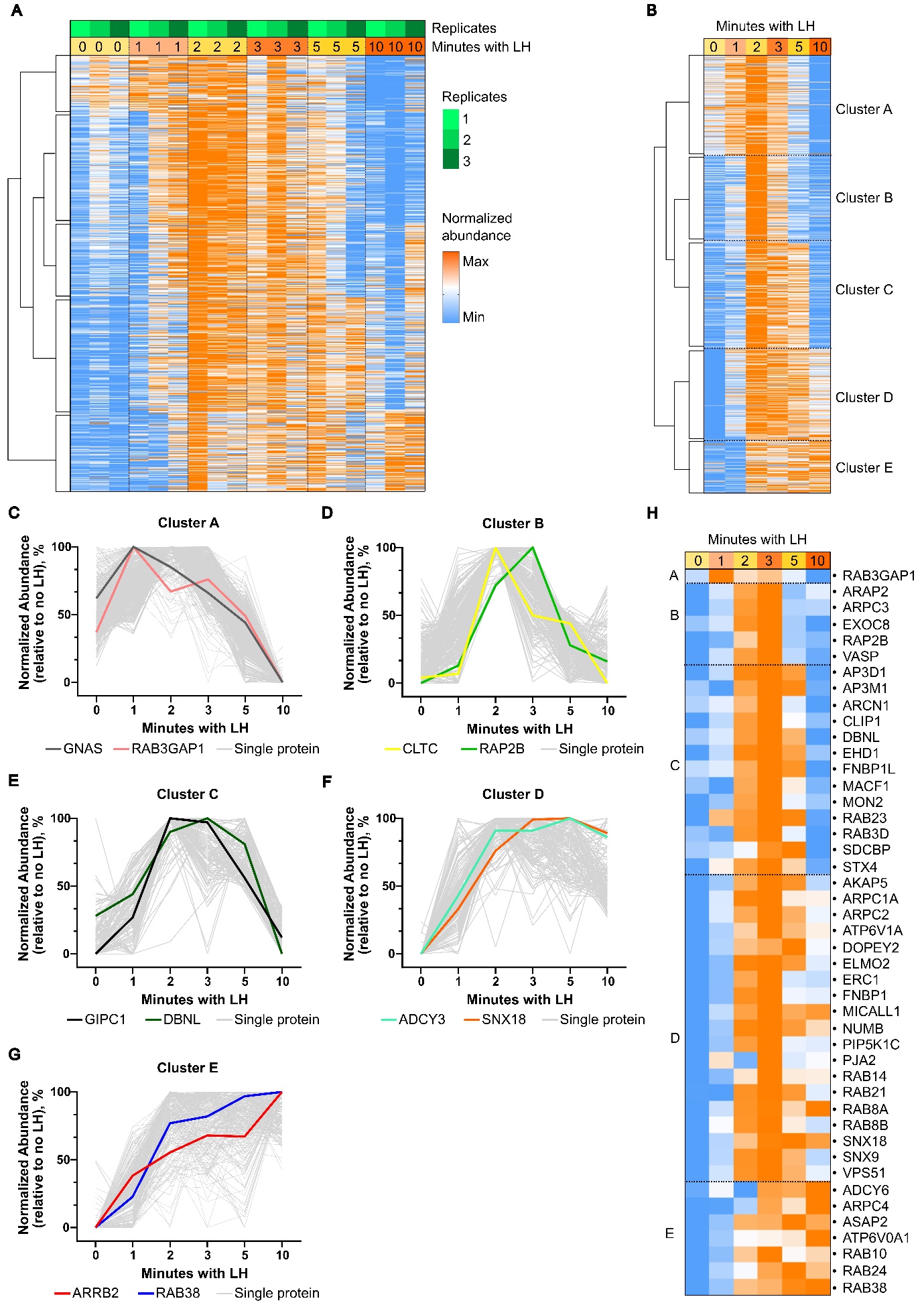

**Figure S6 (relates to Figure 3). A.** Heatmap of hierarchical clustering of interaction profiles of 2439 proteins across three replicates. Columns arranged by stimulation time with LH. Each protein’s abundance across different time points is normalized to “no LH” control first, followed by normalization to its maximum and minimum enrichment signal. Clustering was performed in Perseus using the built-in algorithm. See also **Data S3** for data tables. **B**. Heatmap of hierarchical clustering of averaged interaction profiles of 2439 proteins with five distinct clusters (A-E) shown. Each protein’s abundance across different time points is normalized to “no LH” control first, followed by normalization to its maximum and minimum enrichment signal. Data represents mean (n = 3). **C**-**G**. Representative protein interaction profile plots for each defined cluster. Each cluster plot shows profiles of both a known interactor and a putative hit identified in **Figure 3**, **STEP 4**. Data represents mean (n = 3). **H**. Heatmap of averaged interaction profiles of 45 hits showing their distribution across all five distinct clusters from A to E. Each protein’s abundance across different time points is normalized to “no LH” control first, followed by normalization to its maximum and minimum enrichment signal. Data represents mean (n = 3).

**Figure S7**

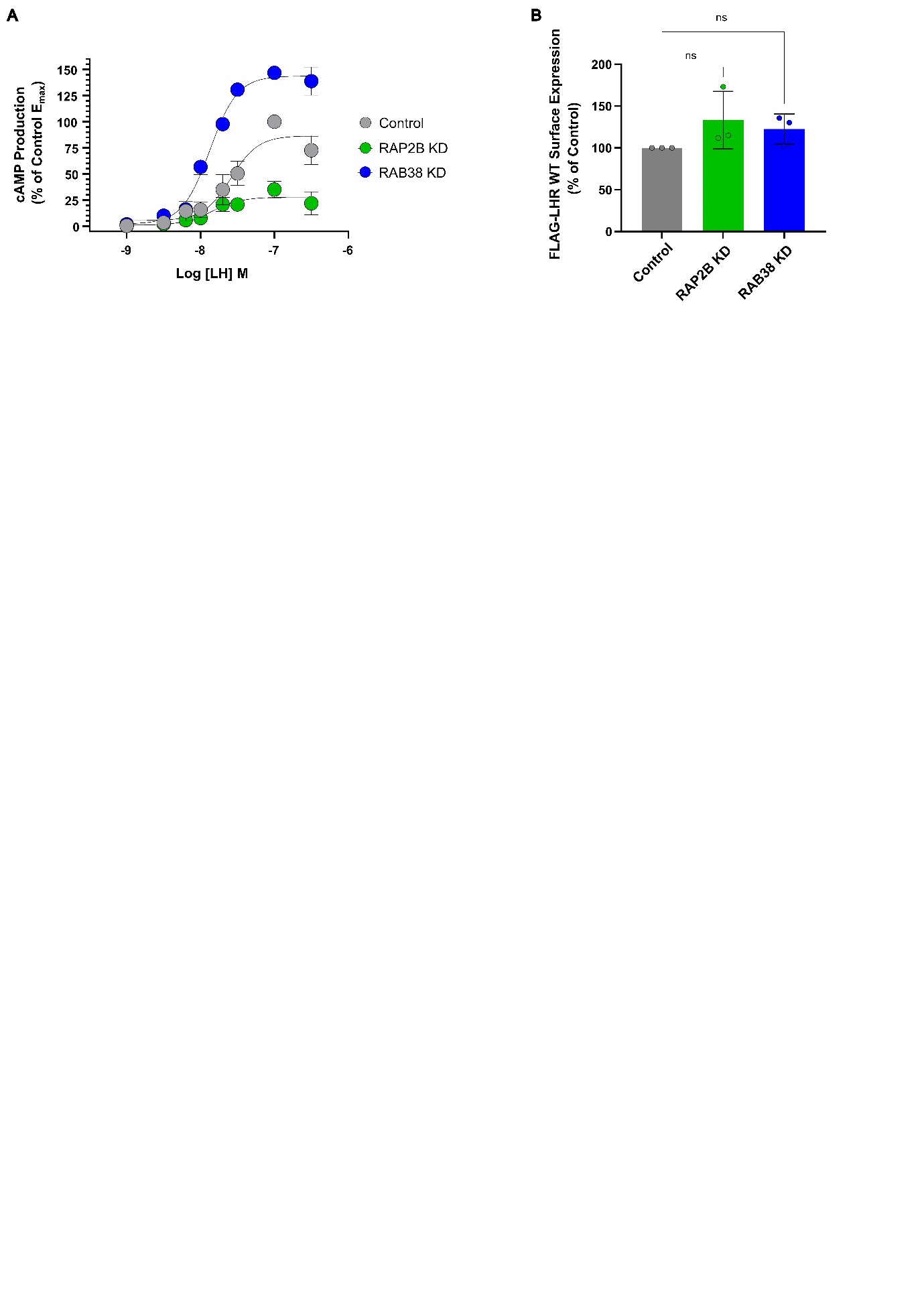

**Figure S7 (relates to Figure 4)**. **A**. LH-induced cAMP concentration-response curves measured by TR-FRET cAMP assay showing the effect of RAB38 and RAP2B knock-downs (KD) on the LHR function. Values were normalized to the NTC (Control) E_max_ (100%). Compared to NTC (Control), RAB38 KD caused an increase in the signaling efficacy (150%) and potency, whereas RAP2B KD decreased the efficacy to 40%. Data represents mean ± SEM (n = 3). **B**. Flow cytometry showing no effect of either RAB38 or RAP2B knock-down (KD) on FLAG-LHR WT cell surface expression. Values were normalized to the NTC (Control) surface expression (100%). Data represents mean ± SEM (n = 3). Statistical analysis was performed by unpaired Student’s t-test: ns > 0.05.
